# Subcellular spatially resolved gene neighborhood networks in single cells

**DOI:** 10.1101/2022.08.03.502409

**Authors:** Zhou Fang, Adam J. Ford, Thomas Hu, Nicholas Zhang, Athanasios Mantalaris, Ahmet F. Coskun

## Abstract

Mesenchymal stem cell (MSC)-based therapies have offered promising treatments against several disorders. However, the clinical efficacy and consistency remain underdeveloped. Single-cell and bulk molecular analyses have provided considerable heterogeneity of MSCs due to origin, expansion, and microenvironment. Image-based cellular omics methods elucidate ultimate variability in stem cell colonies, otherwise masked by bulk omics approaches. Here, we present a spatially resolved Gene Neighborhood Network (spaGNN) method to produce transcriptional density maps and analyze neighboring RNA distributions in single human MSCs and chondrocytes cultured on 2D collagen-coated substrates. This proposed strategy provides cell classification based on subcellular spatial features and gene neighborhood networks. Machine learning-based clustering of resultant data yields subcellular density classes of 20-plex biomarkers containing diverse transcript and protein features. The spaGNN reveals tissue-source-specific MSC transcription and spatial distribution characteristics. Multiplexed spaGNN analysis allows for rapid examination of spatially resolved subcellular features and activities in a broad range of cells used in pre-clinical and clinical research.

## Introduction

Mesenchymal Stem Cells (MSCs) show strong regenerative and immunoregulatory potential, and are tested as cell-based therapies for a wide range of diseases.^1, 2^ Owing to their differentiation potential into clinically relevant lineages such as osteocytes, chondrocytes, and adipocytes and their immunomodulatory role, MSCs are widely studied and have been shown to display considerable spatial and temporal molecular heterogeneity.^3^ MSCs are isolated from multiple sources including human bone marrow (HBM), adipose tissue (HAT), and umbilical cord (HUC), making it challenging to clinically validate the efficacy and safety of cell-based therapies.^2, 4^ Donor age and general health status introduce further MSC heterogeneity.^5^ Allogenic MSCs benefit biomanufacturing processes with wide availability.^6^ However autologous MSCs eliminate the issue of immune rejection associated with allogeneic MSCs.^7^ The standardization of MSC-based therapies is further complicated by the intra-colony variability of MSCs due to clonal evolution during manufacturing.^8^ MSC phenotype is currently defined by the expression of *CD105*, *CD73*, and *CD90* markers, lack of expression of *CD45*, *CD34*, *CD14* or *CD11b*, *CD79a* or *CD19*, and *HLA-DR*, and the potential to differentiate into osteoblasts, adipocytes, and chondrocytes *in vitro.*^9, 10^ However, deep molecular profiles beyond surface markers are required to understand MSC heterogeneity.

Bulk transcriptomics, proteomics, and metabolomics approaches have been used to profile MSCs from distinct donors and tissue sources, but molecular details of individual cells are masked due to population-wide molecule extraction protocols.^11–13^ Single-cell RNA sequencing of human MSCs has yielded functional subpopulation differences in differentiation potential, immunoregulatory function, and clinical efficacy at the cost of subcellular spatial details.^14–18^ Label-free and morphological imaging methods have demonstrated the utility of image-based MSC classification for clinical cell manufacturing pipelines using only a limited number of biomarkers.^19–21^ Spatially resolved molecular omics technologies can combine the advantages of imaging approaches and single-cell molecular analyses while providing more detailed spatial molecular profiling of MSCs.

Emerging spatially resolved and multiplexed gene expression profiling technologies include two mainstream approaches. Spatial transcriptomics and Slide-seq leverage DNA-barcoded substrates and microbeads to capture multiple RNA targets, providing gene expression maps of tissues at 10-100 µm resolutions.^22, 23^ The spatial resolution limitation and sample preparation complexity of these platforms are bottlenecks for transcriptional analysis of single stem cells. Another approach utilizing image-based barcoding of gene expression maps is the Fluorescence In-Situ Hybridization (FISH) method for detection of RNA molecules in their native positions within cells and tissues. Sequential FISH (SeqFISH) and multiplexed-error-robust FISH (MERFISH) methods have generated single-cell spatial maps of 10-10000 RNA species in mouse tissues. These maps have since been utilized in neuroscience, developmental biology, and embryonic stem cell biology^24–27^. While these demonstrations were initially limited to murine models, emerging multiplexed FISH techniques combined with signal amplification assays open doors to the detailed study of human stem cell biology, deciphering subpopulation, donor, and source heterogeneity of MSCs.^28^

The subcellular organization of RNA molecules inside individual cells is crucial to position-dependent transcriptional control of stem cell differentiation and cellular decision-making processes.^29^ Thus, subcellular spatial transcriptomics can provide regulatory and functional cell information otherwise masked by traditional molecular analyses. FISH-based multiplexing methods localize single RNA transcripts, enabling the study of the spatial organization of RNA-enriched compartments in subcellular volumes, a feature previously limited to live-cell imaging.^30^ For instance, *β-Actin* molecules are observed to be localized at the cell edge and focal adhesion points, allowing cells to efficiently migrate and polarize.^31^

In cells, complex regulatory relationships between genes maintain gene-expression phenotypes.^32^ Gene transcription regulation is modulated by a cascade of biochemical processes, leading to control of transcription levels.^33^ Gene regulatory networks (GRNs) have been widely used to describe the regulation of gene expression, and a wide range of tools have been developed to discover the regulatory relationship between genes.^34–37^ GRNs provide intuitive visualization by representing genes as nodes in graphs, and regulatory relationships as edges. Co-expression networks reveal functional and regulatory commonalities between genes.^38, 39^ However, they lack directional connection and miss the regulator or target information. Investigation of transcription factor binding sites in the genome dissects the biochemical process of regulation.^35, 40, 41^ However, regulatory relationships are frequently confounded by multiple regulatory pathways that lead to imprecise models.^34^ Statistical inference models have also been proposed to infer regulatory networks from single-cell transcriptomic profiles.^36, 37^ Spatial transcriptomics data has also been used to infer gene regulatory relationships.^42, 43^ These computational approaches utilize existing transcriptomics data to predict regulatory relationships. Although these expedient methods utilize widely available data, they too lack subcellular spatial resolution. (**Table 1, Table 2**)

**Table 1.** Summary of current gene network analysis methods.

| Computation approach | Summary | Technology and data | Examples |
| --- | --- | --- | --- |
| Co-expression | Discover co-expression and correlation relationships. Does not infer direct regulation relationship. | scRNAseq<br>DNA microarray<sup>38</sup> | Stuart, *et al.* 2003<sup>38</sup><br>Tang, *et al.* 2018<sup>39</sup> |
| Transcription factor binding sites analysis | Directly uses binding site information and biochemical process of regulation to study gene expression regulation.<sup>34</sup><br>Result can be confounded by multiple transcription regulation pathways.<sup>34</sup> | DNA sequencing, sequence matching,<br>ChIP-seq | Daxinger, *et al.* 2021<sup>41</sup><br>Kulkarni, *et al.* 2018<sup>35</sup><br>Verfaillie, *et al.* 2015<sup>40</sup> |
| Computational inference | Infers regulatory network from gene expression data using mathematical models. | scRNAseq | Liu, *et al.*, 2022<sup>36</sup><br>Huynh-Thu, *et al.* 2010<sup>37</sup> |
| Spatial regulation | Infers regulatory relation<br>from spatially resolved<br>transcriptomics using<br>spatial patterns | FISH, spatial<br>transcriptomics | Yang, *et al.*, 2019<sup>42</sup><br>Puniyani and Xing,<br>2013<sup>43</sup> |
| SpaGNN (The<br>proposed method) | Examines cellular,<br>subcellular, and localized<br>co-expression | Image-based<br>spatial<br>transcriptomics |  |

**Table 2.** Selection considerations of current gene network analysis methods and spaGNN.

| Computation approach | Advantage | Disadvantage |
| --- | --- | --- |
| Co-expression | Computationally identified using a straightforward metric. | Lacks inference of regulator and target genes. Connection is not directional. |
| Transcription factor binding sites analysis | Directly studies the regulator of target genes. Regulatory relationships are more likely to be realistic. | Regulatory relationship can be weak due to multiple regulators controlling the same target.<sup>34</sup> |
| Computational inference | Data widely available. Can analyze highly multiplexed data. | Computationally complex. Prediction ability varies between different models and datasets.<sup>85</sup> |
| Spatial regulation | Utilizes spatial distribution of RNA molecules. | Loss of spatial variation data of regulatory networks. |
| SpaGNN (The proposed method) | Examines cellular, and subcellular colocalization and correlation.<br>Examines variation of subcellular RNA spatial organization. | Limited to colocalization and correlation study. |

Here we present a new analytical method, spatially resolved gene neighborhood network (spaGNN), that constructs subcellular molecular density maps to distinguish cell state differences of individual MSCs. This pipeline acquires multiplexed, spatially resolved subcellular RNA maps of single cells through sequential and multiplexed Hybridization Chain Reaction (HCR), a robust amplification strategy utilized in the detection of single RNA molecules present in human specimens (**Fig. 1**). For proof-of-concept, we developed an analysis pipeline to process the multiplexed data for generating “gene neighborhood” maps in HBMs, HUCs, and human chondrocytes (HCHs). HCH was included as a clinically relevant lineage of MSCs due to the wide application of MSC-based therapies for cartilage regeneration.^44, 45^ First, a machine-learning-based clustering algorithm of multiplex data was used to classify regions in subcellular volumes of individual MSCs termed “RNA enrichment regions.” Subsequent protein staining validated the structural features and protein distribution of the cell. Later, subcellular “patches” were then defined by local RNA density or subcellular spatial regions. The correlations of RNA species among patches and neighbors were then analyzed to reveal the subcellular organization of RNA. Thus, the presented spaGNN workflow decodes cellular and subcellular transcriptional regulation of MSCs from distinct donors and sources, providing many opportunities to define the identities of MSC subpopulations.

**Figure 1.**
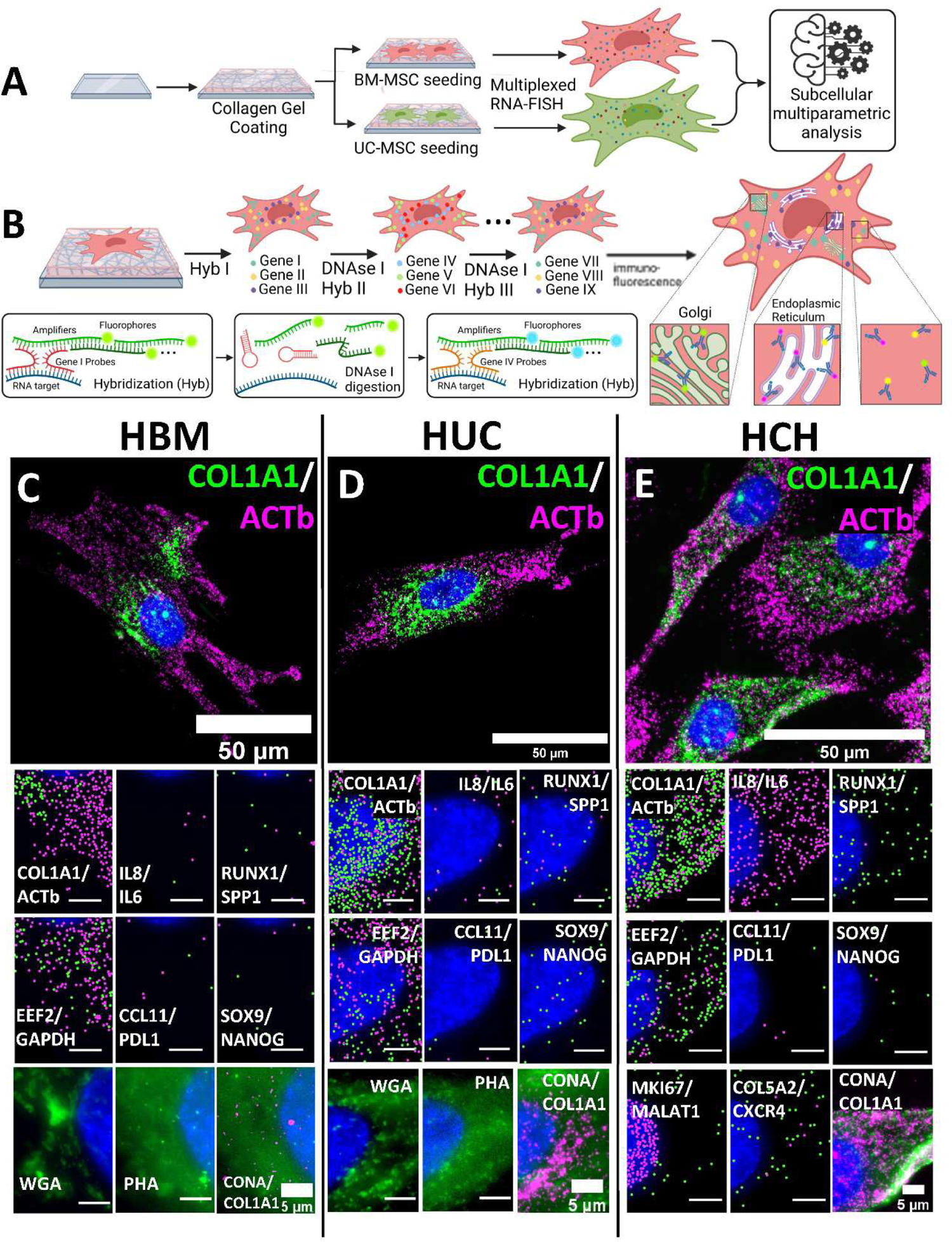
Multiplexed HCR-RNA method profiles MSC spatial transcriptomics at single RNA resolution. **(A)** Glass coverslips coated with collagen gel enable cell adherence. Cell types of interest, namely HUC (n=121), HBM (n=237) and HCHs (n=247), were seeded, fixed, and processed via multiplexed RNA-FISH and subcellular multiparametric analysis. **(B)** Fixed cells were hybridized, digested, and rehybridized for all gene targets. Hybridization occurs via self-assembly of DNA probes designed to attach up and downstream of an RNA target. Probes conjugate with a series of amplifiers joined to a fluorophore in a process called Hybridization Chain Reaction (HCR). These markers are then digested with a DNase I enzyme and new markers can be placed for a different target. After all targets are identified, Immunofluorescence staining is performed for protein visualization and organelle identification. **(C)** Multiplexed RNA profiles of 12 genes and 3 cellular structure markers on Human Bone Marrow derived MSC. **(D)** Multiplexed RNA profiles of 12 genes and 3 cellular structure markers on Human Umbilical Cord derived MSC. **(E)** Multiplexed RNA profiles of 16 genes and 3 cellular structure markers on Human Chondrocytes. Scale bars without a label have a length of 5µm.

## Results

### Spatially resolved transcriptional and proteomic profiling of stem cells

Human MSCs exhibit transcriptional heterogeneity at the single-cell, tissue source, and donor levels. HUC-MSCs, for example, show distinct differences in functional characteristics such as proliferation potential, differentiation efficiency, and immunomodulatory character when compared to HBM-MSCs.^46^ Autologous HBM-MSCs reduce inflammation in regenerative therapies, while allogeneic sources demonstrate efficacy in only a subset of patients.^47^ We reasoned that allogeneic HBM-MSCs and HUC-MSCs would be a crucial proof-of-concept demonstration for studying spatially regulated transcriptional diversity of tissue-specific MSCs due to their prevalent use in current clinical and pre-clinical research. Utilizing the spaGNN workflow, we examined HUC-MSCs (n=121), HBM-MSCs (n=237) and HCHs (n=247).

MSC samples were obtained from RoosterBio based on clinical quality-control standards (**Table S1**) to ensure the studied cell populations exhibited similar bulk characteristics to the therapeutic product. To overcome the imaging limitations of traditional cell culture methods on plastic substrates, we cultured cells at a density of 500 cells/cm^2^ on an optically clear 2D glass substrate (100-150 µm thickness) coated with a sub-5-micron thick layer of collagen. Previously, multiplexing of RNA labeling using seqFISH protocols was achieved by sequential hybridizations, imaging, and re-labeling of 10-10000 RNA species.^24, 48, 49^ However, MSCs are structurally intricate stromal cells exhibiting a high cell autofluorescence background due to high subcellular protein density. Therefore, spaGNN analysis of MSCs necessitates the use of the split-probe design and signal amplification of the HCR assay^28^, providing bright, single-molecule RNA images with high detection specificity (**Fig. 1A**). This multiplexing strategy simultaneously labels and images up to three RNA targets per cycle. Each cycle of HCR and imaging is followed by enzymatic digestion of HCR components using a DNase I enzyme. Subsequent rounds of labeling then target different sets of RNA species, providing a linearly scalable method of cycle-number dependent multiplexing. For example, six sequential hybridizations with three RNA targets per hybridization would profile eighteen unique RNA markers in a single MSC (**Fig. 1B**).

Next, multiplexed RNA maps in single MSCs were combined with image-based morphological analysis. The spaGNN workflow implements a cell painting strategy to co-detect cellular structures in the same MSC.^50^ After completing the multiplexed RNA imaging, the sample is stained with Concanavalin A (*ConA*, staining Endoplasmic reticulum), Phalloidin (*PHA*, staining Actin), and Wheat Germ Agglutinin (*WGA*, staining Golgi apparatus and plasma membrane) to visualize subcellular features of each MSC. This method provides cell segmentation reference, and the resultant spatially resolved multi-omics data provides subcellular transcriptional profiles and structures of HUC, HBM, and HCH.

Spatially resolved gene expression maps of single MSCs were provided in the form of individual images for multiple RNA targets and cell-painting markers (**Fig. 1C-E**). Observed MSC subcellular heterogeneity suggests region-dependent RNA densities within individual MSCs, a finding contrary to models of RNA distribution utilizing an assumption of Poisson distribution of RNA within a cell. This observation is akin to chromatin density detected in stem cell differentiation.^51, 52^ Such RNA densities have previously been observed in neuronal cells^53^, but not in MSCs.

12 RNA markers were measured in HBM and HUC, and 4 additional RNA markers were measured in HCHs (**Fig. 1C-E, S1-S4**). *NANOG*, a pluripotency marker, was detected in only a few RNA copies per cell and was predominantly clustered around the nuclear membrane. *PDL1*, *IL6*, *IL8*, and *CCL11* are genes related to the immunomodulatory function of MSCs which exhibited differences in their locations relative to cytoplasmic membranes in this experiment. Subsequent cell painting stains demonstrated cell-source specific RNA densities and Golgi apparatus locations, most likely due to ER-enrichment of translational machinery and rapid turn-over of collagen secretions through ER-Golgi transport^26, 54^. *COL1A1* transcripts were observed to be enriched on ER structures as detected by *ConA* staining and were excluded from the Golgi region defined by the *WGA* stains. Another important observation is that neighboring cells exhibited asymmetric RNA distributions, especially *SPP1* and *β-actin* genes in pairs of HUC. This finding can likely be attributed to asymmetric cell division events leading to a unique differentiation potential for each MSC. The initial observations were further investigated using the spaGNN pipeline.

### Clustering analysis of multiplexed subcellular RNA and cell-painting images

Spatially resolved single-cell data has been used to define cell types and their locations in mouse and human tissues^55, 56^. The initial approach will assess subcellular RNA heterogeneity to identify unique spatial regulation of individual RNA segments in human MSCs. Previous work has developed analytical tools to annotate cell types in tissue datasets. These tools use machine learning algorithms based on high-dimensional data clustering and visualization algorithms.^57–59^ However, these tools lack the subcellular spatial specificity capabilities that are demonstrated in the spaGNN pipeline.

The initial images were processed and cross-registered to align cell positions. The Pixels of multiple HBM, HUC, and HCH images were combined to perform clustering of gene-enrichment regions (**Fig. 2A, 2B**). Each cell contains millions of pixels, corresponding to several million data points. To address the large amount of data, pixels were sampled using a geometric sketching algorithm, then down-sampled to 5% of the original size and clustered to represent the distribution of pixel data^60^ (**Fig. 2A**). The Leiden clustering algorithm finds gene enrichment clusters from the subset of all pixels.^61^ To assign each pixel to a cluster, we employed a nearest neighbor search method. For each pixel in the down-sampled image, 30 nearest neighbor pixels in the data feature space were examined, and the pixel was assigned the most likely neighbor label. The blue-red colored heatmap shows the intensity level of each pixel involved in the clustering analysis (**Fig. 2C**). Cell images were then recolored after up-sampling based on the cluster assignment of each pixel previously determined in the down-sampled data set to visualize gene enrichment regions (**Fig. 2A, 2D**).

**Figure 2.**
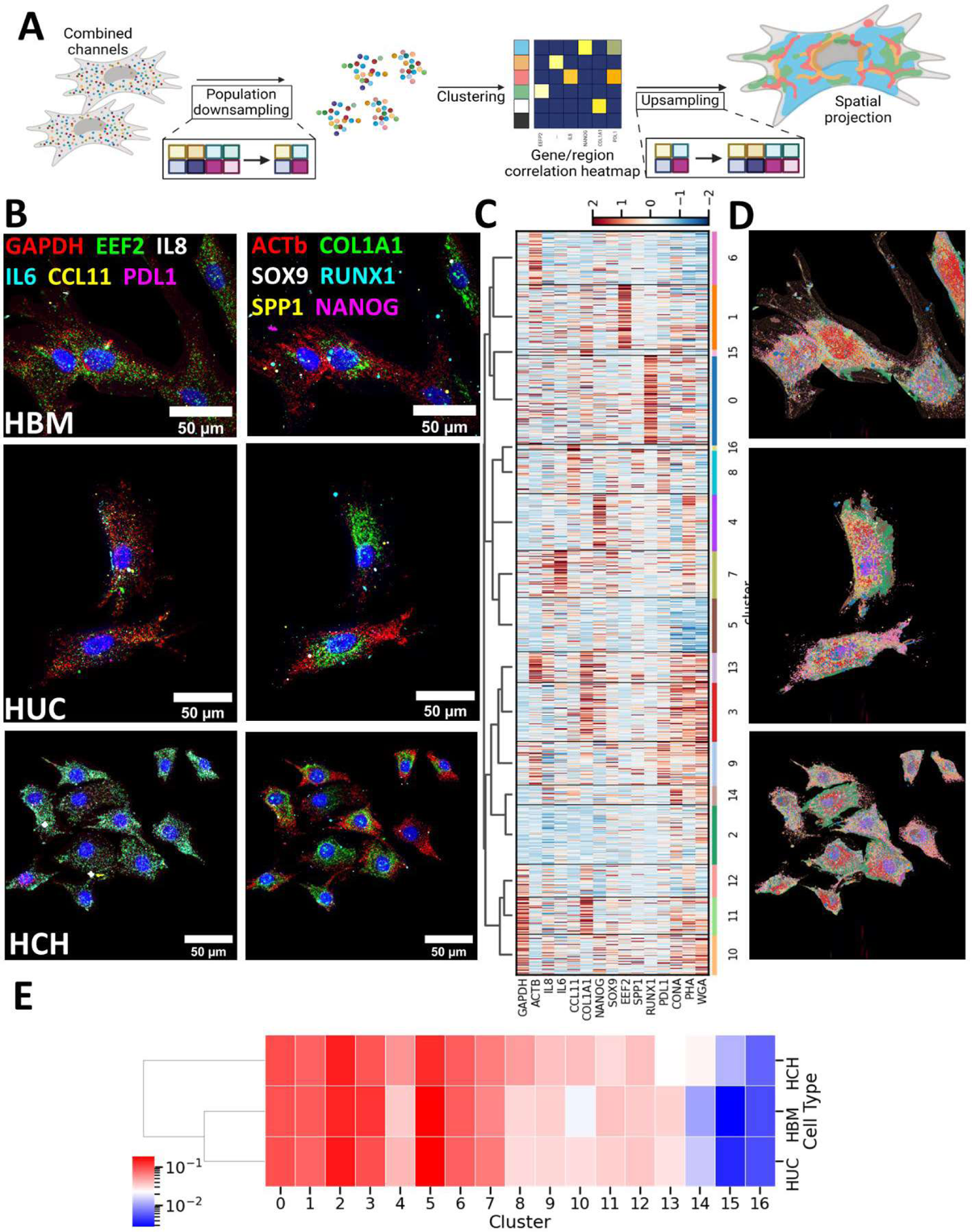
Subcellular clustering reveals spatial gradient patterns in single-cell transcriptomic profiles. **(A)** Illustration of subcellular pixel-level clustering analysis. Pixels were first down-sampled to reduce data size. High-dimensional data for each pixel was clustered using the Leiden algorithm. All pixels were then assigned to a cluster based on the clustering result on the down-sampled data. Cells were then pseudo-colored based on cluster labels of each pixel to reveal spatial distribution of each cluster. **(B)** The registered images of RNA in cells. A phase cross correlation registration method was used to correct for both translational and rotational shifts in images between cycles. The cross-registered images show alignment between images of all markers. **(C)** Heat map of clustered pixels from HUCs (n=121), HBMs (n=237) and HCHs (n=247). All pixels involved in clustering are shown in the heatmap. Blue-red colors indicate the enhancement of markers for the corresponding pixels. Tendency of enrichment of pixels within the same cluster indicates the co-existence of markers in clusters. **(D)** Pseudo-colored cell images according to the cluster assignment of each pixel. All three cell types exhibit gradient of cluster changes from the center of cells towards the cell membrane. **(E)** Heatmap of differential enrichment of each cluster in each cell type. The proportion of pixels belonging to each cluster was computed for each cell type. Enrichment of clusters 0-3, 5-7, 12, and 16 is common to all cell types. HBM shows suppression of clusters 4 and 10. HCH shows suppression of clusters 13, 14, and 15, and enrichment of clusters 8, 9, and 10. HUC demonstrates moderate enrichment of non-common clusters and suppression of clusters 14 and 15 similar to HBM.

The recolored cell images validate that *COL1A1* mRNA is more abundant near the nucleus, while *ACTb* is more abundant in the cytosolic region. Other mRNA molecules are mostly distributed throughout the cell. When comparing resulting pseudo-colored cell images of different cell types, despite tissue source differences, similar subcellular, non-uniform distributions were obtained in HBM, HUC, and HCH (**Fig. 2D, S5**). These findings suggest that the position-dependent transcriptional processes for these gene targets may be universally conserved based on either subcellular structural similarity or highly conserved translational mechanisms. To analyze the enrichment of each cluster in each cell type, the proportion of pixel of a cluster among all pixels were computed for each cell type (**Fig. 2E**). The blue-red colored heatmap indicates the enrichment of each cluster. The dendrogram indicates HBM and HUC are similar in enrichment of clusters, and HCH is different compared to HBM and HUC.

The image-based spaGNN workflow for subcellular clustering and heterogeneous RNA enrichment enables the study of gene regulation and position-dependent transcriptional alterations in stem-cell biology beyond MSCs. Using the scalable and multimodal multiplexing strategy presented herein, complementary transcriptional and proteomic targets may shed light on subcellular mechanisms of Spatio-temporal RNA anchoring on proteomic and organelle structures.^62^

### Statistical comparison of subcellular distribution of RNA

Due to the observed centralized *COL1A1* distribution, we analyzed subcellular enrichment of RNA as a function of distance from the center of the cell (**Fig. 3**). Cell centers were defined as the center of mass of the cell segmentation mask. The number of RNA signals as a function of distance from the cell center showed similar patterns between different RNA markers, as most RNA molecules analyzed in this study indicated cytosolic localization. However, different RNA signals exhibited maximum enrichment at different distances. For example, the peak expression of *COL1A1* was closer to the center of the cell than the peak expression of *EEF2* (**Fig. 3A**). Visualization of RNA molecule positions and their distances from the cell center also demonstrated that *COL1A1* congregates about the perinuclear region (**Fig. 3B**). The same analysis was expanded to all HBM, HUCs, and HCHs; enrichment of each gene versus distance from cell center was visualized as a heatmap (**Fig. 3C**). The subcellular distribution of RNA of different cells was then analyzed using a Kolmogorov–Smirnov Hypothesis Test, yielding cell type comparisons. The p-values are visualized in a heatmap (**Fig. 3D**).

**Figure 3.**
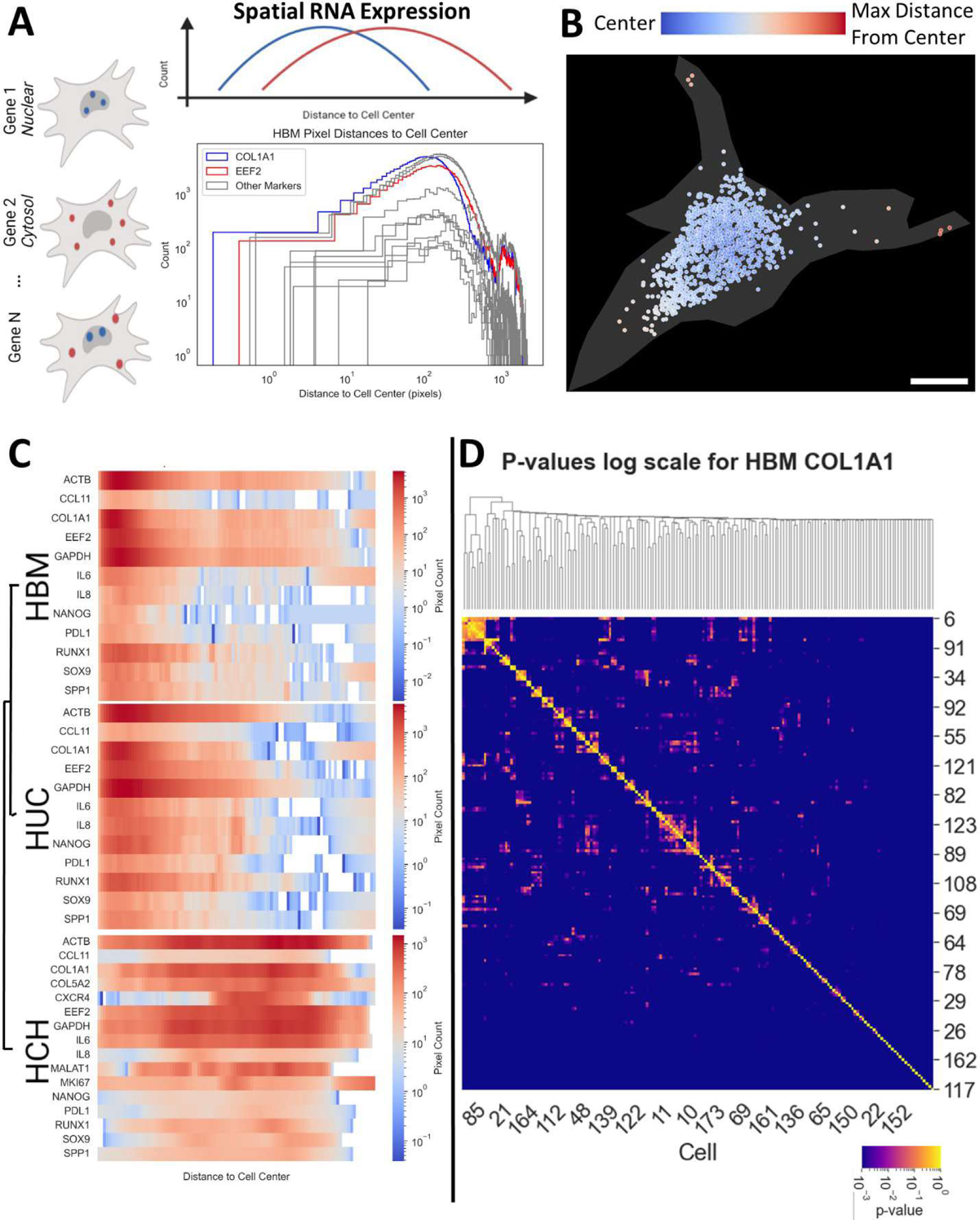
Statistical comparisons of cell source populations confirm distinct spatial patterning of RNAs. **(A)** Spatial RNA expression measured with respect to each cell’s center of mass. RNA molecules are nuclear or cytosol associated as illustrated in the histogram of each transcript’s distance to the cell center. Examples highlighted in blue and red are *COL1A1* and *EEF2* genes respectively in a single HBM. **(B)** Visual validation of spatial RNA pattern (blue/red) on cell mask (gray). Example shown is *COL1A1* in an HBM. Blue dots indicate shorter distance from the center of the cell and red dots indicate longer distance from center of the cell. The scale bar indicates a length of 10µm. **(C)** Spatial RNA histograms as heatmaps for each gene for HUCs (n=121), HBMs (n=237) and HCHs (n=247). HCH genes exhibit more cytosolic patterns than corresponding genes for HBM and HUC, while HUC and HBM present higher similarity in spatial distribution of RNA. **(D)** The graph shows the p-values of the Kolmogorov–Smirnov Hypothesis Test of *COL1A1* spatial relationships for all pairwise cells in the HBM population.

### Subcellular spatial patch analysis

Previous analyses have established that subcellular distributions of RNA are not uniform. Subcellular, pixel level clustering results indicate various RNA enrichment regions; enrichment of RNA with respect to the center mass of the examined cells showed varying subcellular spatial distribution of RNA. Therefore, we investigated the separability of each cell into patches defined by high local RNA density utilizing the Leiden clustering algorithm to detect similar data points^63^ (**Fig. 4A**). When applied to positions of all detected transcripts, the algorithm separates subcellular patches by local RNA density peaks. Pearson’s correlation between markers among patches indicates the tendency for transcripts to colocalize in the same patch (**Fig. 4A, S6**).

**Figure 4.**
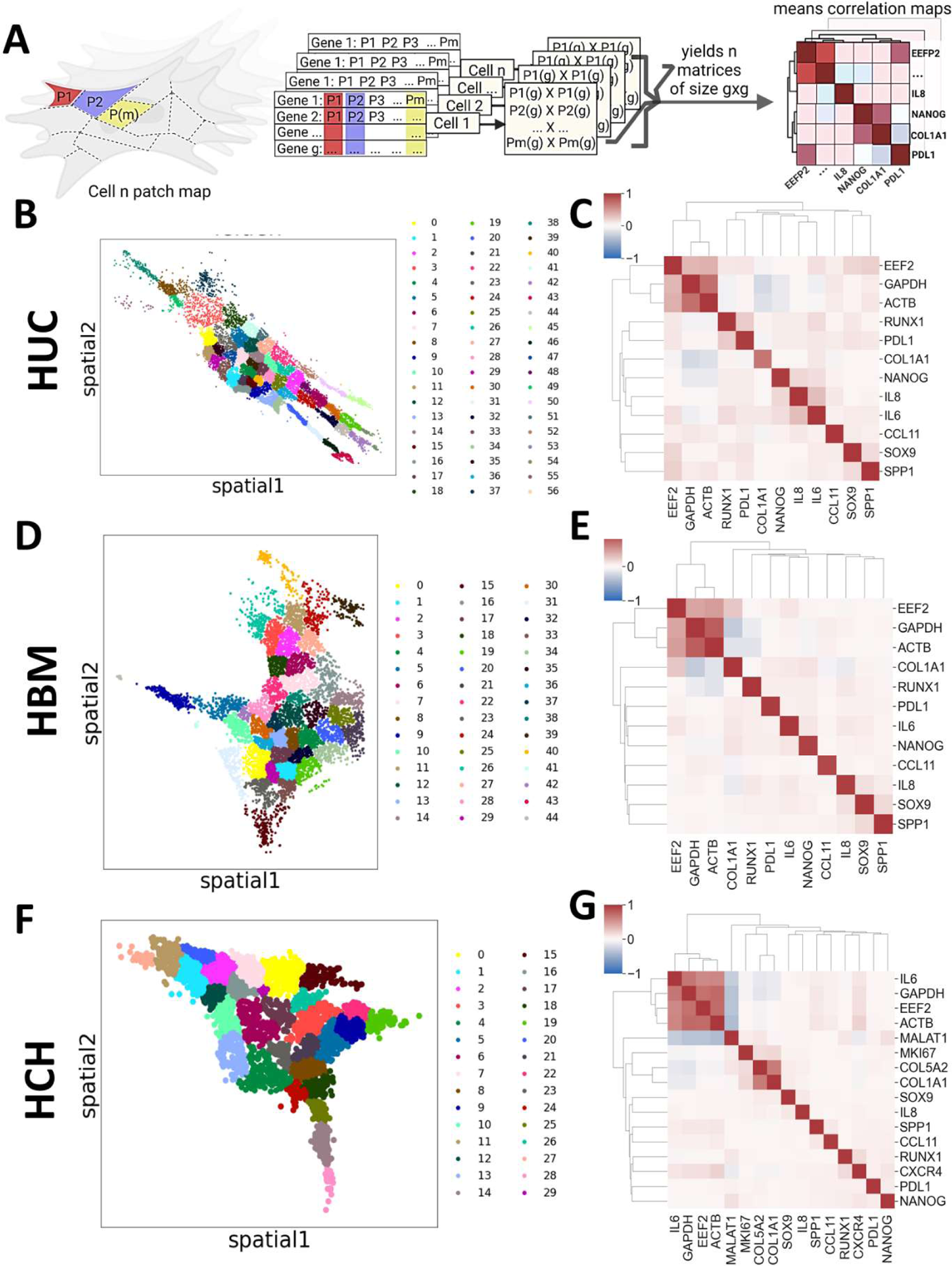
Patch level analysis reveals subcellular heterogeneity in spatial transcriptomic profiles. **(A)** Local high RNA density regions were detected using the Leiden clustering algorithm. Each species of RNA (g) was counted, and correlations of RNA count between all (m) patches of the same cell (n) was calculated. The process was repeated for all n, generating n means correlation maps of size g x g. **(B)** Leiden clustering of positions of RNA molecules in a HUC cell. The presented cell contains 57 patches. **(C)** Mean correlation map of HUCs (n=121). *COL1A1* shows negative correlations with *GAPDH* and *ACTb*. *EEF2* shows positive correlations with *GADH* and *ACTb*. **(D)** Leiden clustering of positions of RNA molecules in an HCH cell. The presented cell contains 30 patches. **(E)** Mean correlation map of HBMs (n=237). In addition to *COL1A1*, *RUNX1* also shows negative correlations with *GAPDH* and *ACTb*. **(F)** Leiden clustering of positions of RNA molecules in an HCH cell. The presented cell contains 56 patches. **(G)** Mean correlation map of HCHs (n=247). The studied HCH population show distinct correlation features. *IL6* is highly correlated with *GAPDH*, *EEF2*, and *ACTb*. The addition of nucleus concentrated *MALAT1* marker also shows negative correlations with cytosol-associated markers such as *GAPDH*, *EEF2*, *ACTb*, and *IL6*.

The average patch correlation of HBM (**Fig. 4B, 4C**), HUC (**Fig. 4D, 4E**), and HCH (**Fig. 4F, 4G**) are shown. Notable differences include greater *IL6* and *IL8* colocalization in HUC when compared to HBM. Additionally, *IL6* was more colocalized with *GAPDH*, *EEF2*, and *β-actin* in HCHs than HBM and HUC sources (**Fig. 4C, 4E, 4G**). To assist in identifying these differences, a principal component analysis (PCA) was conducted. The original data points were projected to orthogonal axes known as principal components (PCs) to capture the maximum variance in the data while minimizing the number of orthogonal principal components.^64^ PC1-PC2 plotting indicates that HCHs differ from MSCs along PC1, while HBM and HUC cells exhibit more separation along PC2 (**Fig. 5A, S7**).

**Figure 5.**
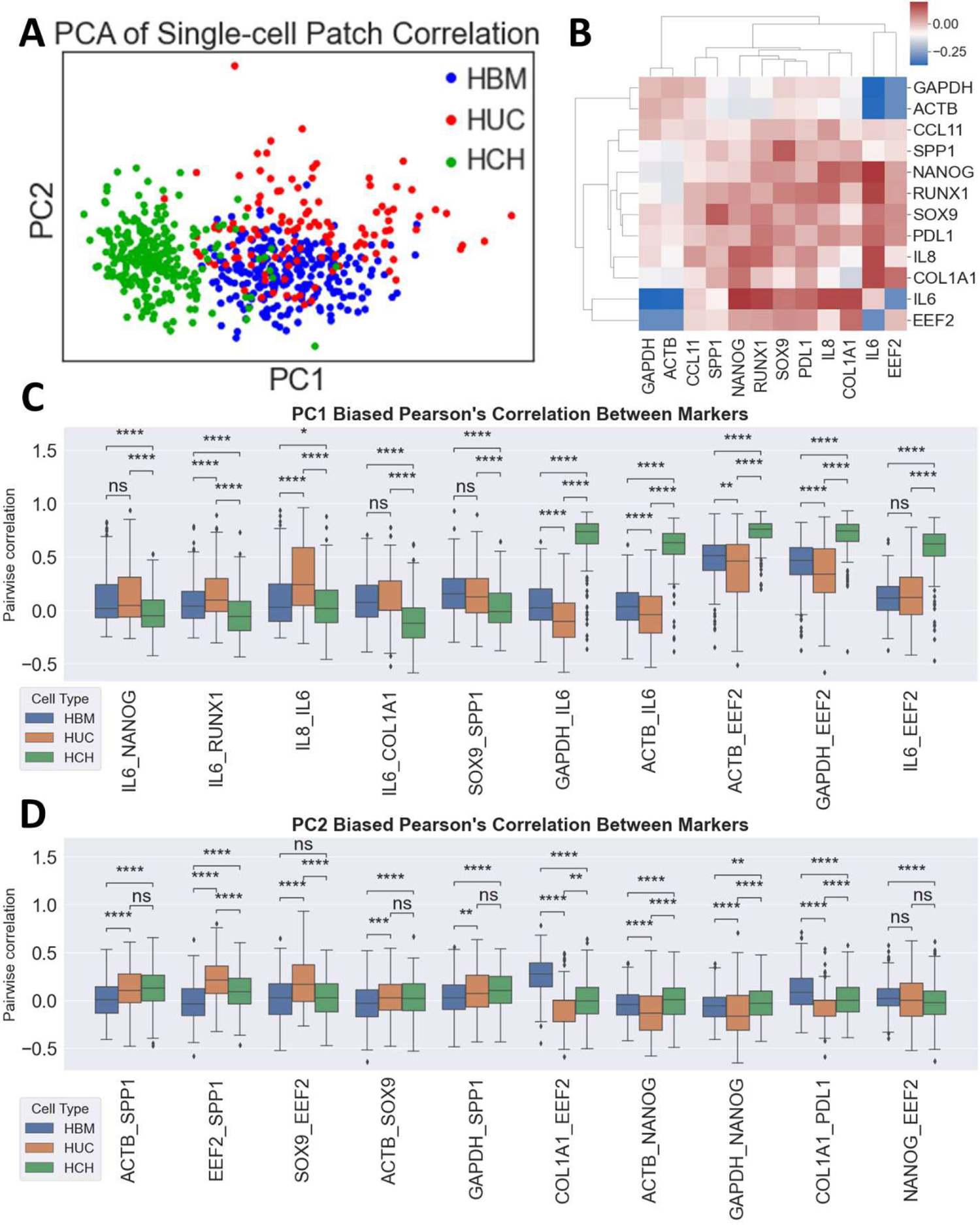
Principal component analysis detects statistically significant features between HBM, HUC, and HCH cells. **(A)** PC1-PC2 plot of patch-level correlation. The plot shows overlap of HBM and HUC, with HCH cells displaying little to no overlap. Separation of HCH and MSCs occurs primarily along PC1. Separation of HBM and HUC is less apparent but appears along PC2. **(B)** Loading of each pairwise correlation in PC1 visualized in blue-red color-coded heatmap. The blue color indicates the correlation is negatively biased in PC1. The red color indicates the correlation is positively biased in PC1. **(C)** Statistical comparison of PC1 biased correlations between HBM, HUC, and HCH. MSCs and HCH are distinguished along PC1. Mann-Whitney-Wilcoxon tests were conducted to examine the differences in pair-wise correlation between cells. **(D)** Statistical comparison of PC2 biased correlations between HBM, HUC, and HCH. MSCs and HCH are distinguished along PC2. Mann-Whitney-Wilcoxon tests were conducted to examine the differences in pair-wise correlation between cells. P-value annotation: ns: 5.00e-02 < p <= 1.00e+00; *: 1.00e-02 < p <= 5.00e-02; **: 1.00e-03 < p <= 1.00e-02; ***: 1.00e-04 < p <= 1.00e-03; ****: p <= 1.00e-04.

The loading of PC1 shows the relation between PC1 and patch correlations (**Fig. 5B**). High PC1 loadings indicate higher correlations, leading to a larger PC1 value. By analyzing features with the highest loadings in PC1 and PC2, we were able to identify significantly different patch correlations between cell types. Statistical analysis of patch correlations with the top 5 positive and negative loadings in PC1 and PC2 was conducted via a Mann-Whitney-Wilcoxon test to compare differences between HBM, HUC, and HCH (**Fig. 5C, 5D**). In correlations with large PC1 loading, several features fail to show significant differences between HBM and HUC cells. However, all compared HCHs and MSCs show statistically significant differences, except for *IL8* and *IL6* differences between HBM and HCHs. Noticeably, most differences between HCHs and MSCs were correlations involving *IL6*, such as *IL6* and *NANOG*, *IL6* and *GAPDH*, and *IL6* and *EEF2* (**Fig. 5D, S7D**).

### Subcellular gene neighborhood network analysis

Patch correlation indicates spatial colocalization at a subcellular patch level, and uncovers differences between HBM, HUC, and HCH cells. Further analysis of localized spatial gene neighbor relationships was conducted through local neighborhood analysis. Within each patch, the center transcript and its surrounding 9 neighbors form a 10-transcript neighborhood as shown by connected dots (**Fig. 6A**). Pearson’s correlation was used as an indication of a transcript’s tendency to colocalize in the same neighborhood. The correlation between the two genes is defined as the connectivity of the 2 markers. The correlation relationship is visualized in network format with node size indicating enrichment of the marker in the patch and edge colors indicating connectivity (**Fig. 6A**).

**Figure 6.**
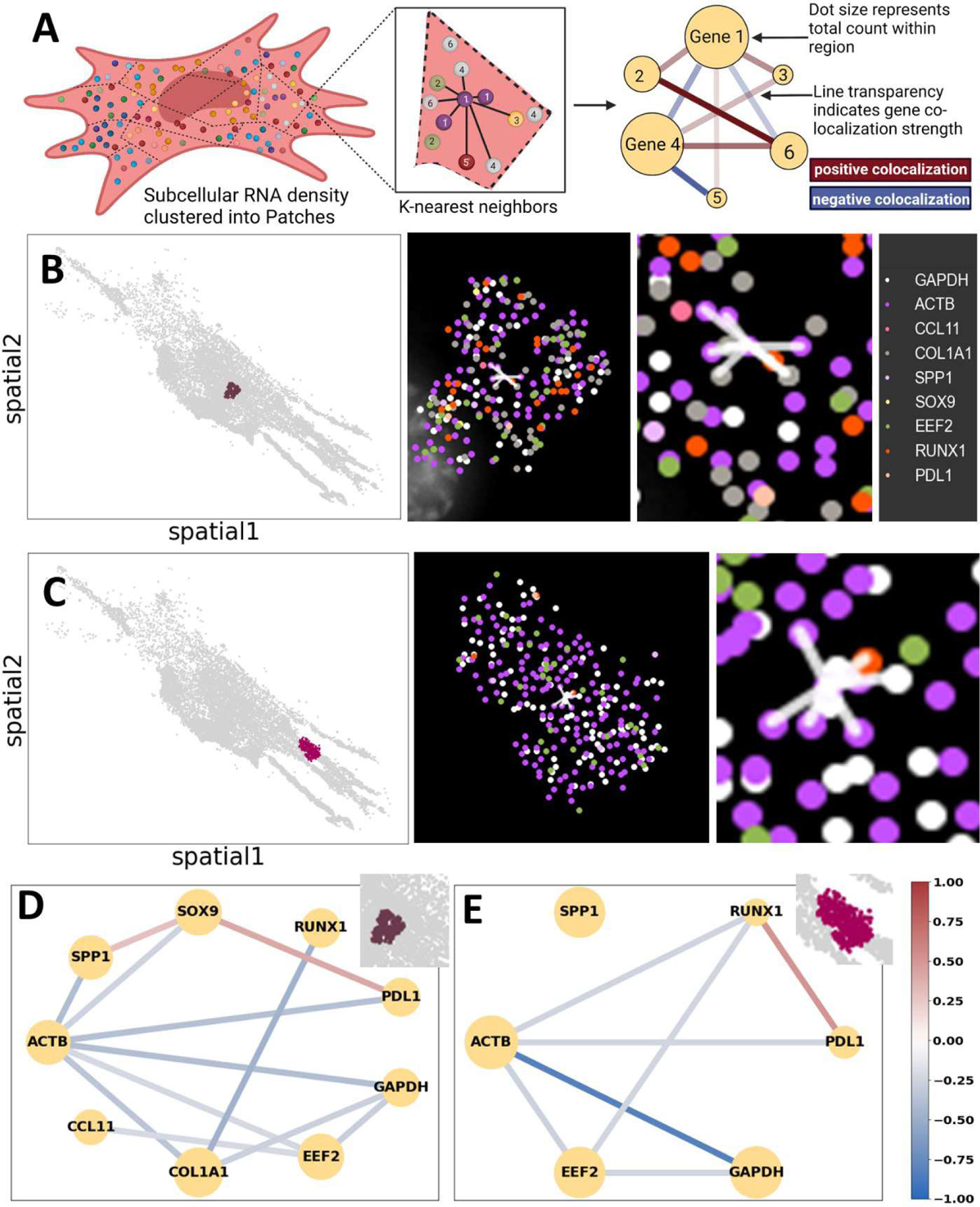
SpaGNN shows subcellular and spatially resolved gene neighborhood networks. **(A)** Subcellular neighborhood analysis pipeline. Center target and its K-nearest neighbors are labeled as a single local neighborhood. Correlation of markers across all local neighborhoods indicates the likelihood of markers to colocalize in the same local neighborhood. Neighboring relationships were visualized as networks per patch. Nodes represent RNA and size of nodes indicates z-score of RNA count in each patch. Edges indicate connectivity of two markers. **(B)** Local neighborhoods in a patch near the nucleus. Scatter plot indicates RNA location in the patch. A local neighborhood is highlighted by connection between center RNA and its neighbors. **(C)** Local neighborhoods in a patch away from nucleus. The patch shows low concentration of *COL1A1*. **(D)** Neighborhood network of the patch indicated in **B**. **(E)** Neighborhood network of a different patch presented in **C**.

The first example demonstrates a patch near the center of the cell, displaying *COL1A1* enrichment (**Fig. 6B, 6D**). The presented neighborhood includes 5 copies of *β-actin*, 1 copy of *GAPDH*, 3 copies of *COL1A1*, and 1 copy of *RUNX1*. The second example demonstrates a patch in the cytosolic region and shows no *COL1A1* expression (**Fig. 6C, 6E**). The neighborhood shown included 6 copies of *β-actin*, 3 copies of *GAPDH*, and 1 copy of *RUNX1*. The same neighborhood connectivity analysis was conducted for all patches among all cells. Across the same cell, patches exhibit varying neighborhood network features (**Fig. 7A, 7B, 7C, S8-S10**).

**Figure 7.**
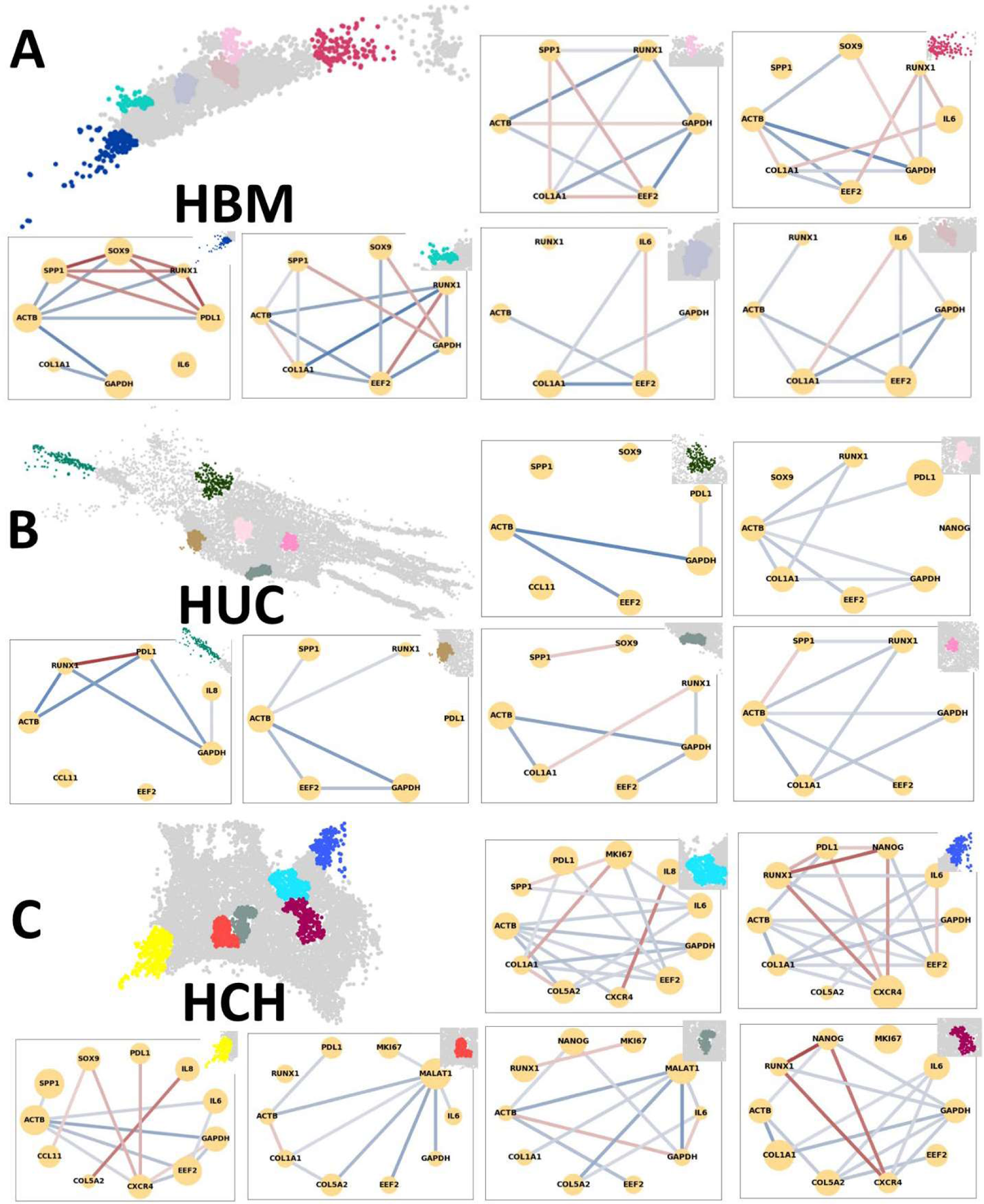
Distinct cell types exhibit heterogeneous gene neighborhood networks Gene neighborhood networks of an HBM (A), a HUC (B), and an HCH (C). Cells present heterogeneous marker enrichment and connectivity between patches. Same marker pairs exhibit different connections in different patches.

**Figure 8.**
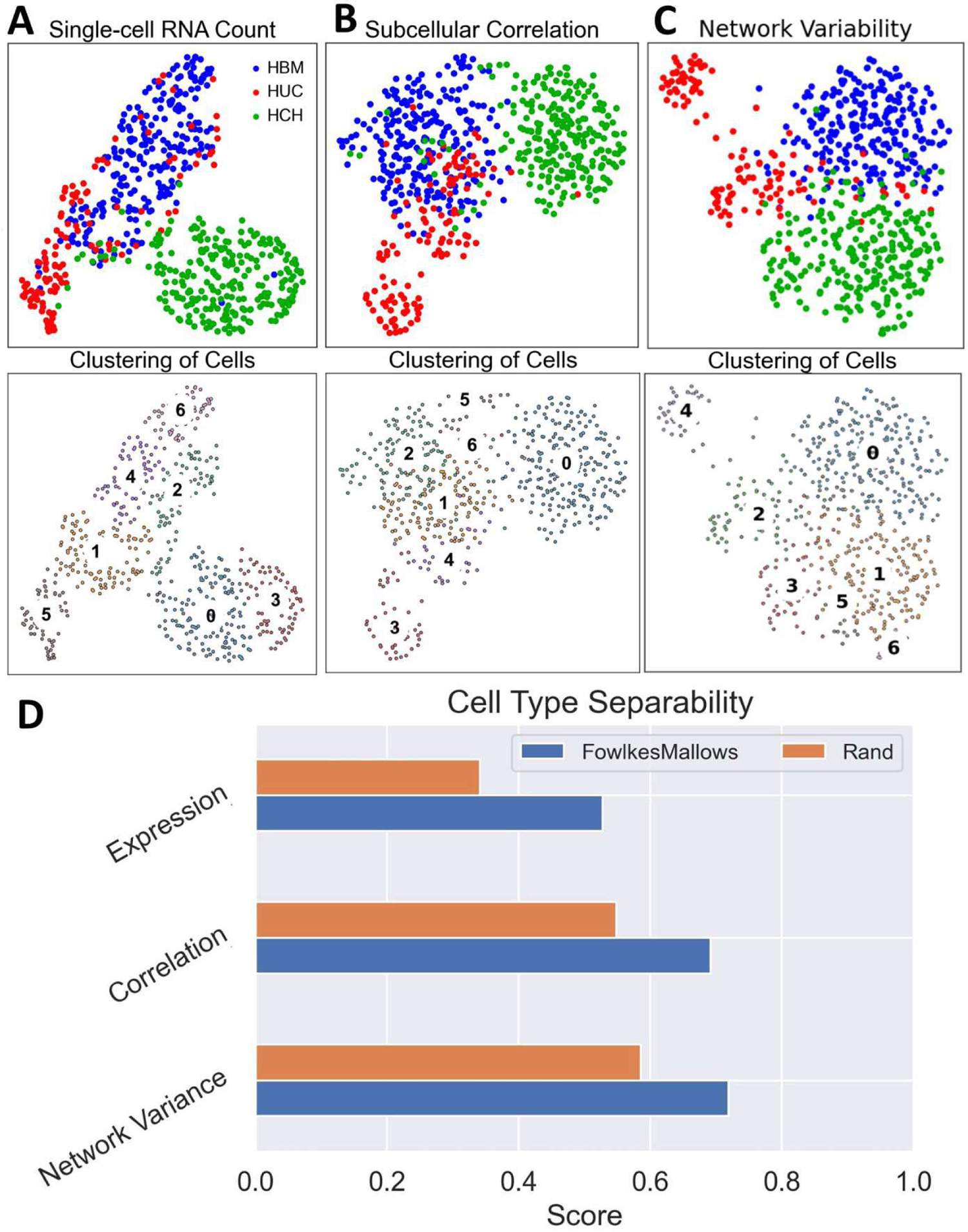
Patch correlation and gene neighborhood network achieve higher separability than single-cell RNA count to distinguish HBM, HUC, and HCH cell populations. **(A)** Cell clustering and cell types based on single-cell RNA count visualized on a UMAP. Count of each RNA type per cell was used as features for UMAP and clustering of the HUCs (n=121), HBMs (n=237), and HCHs (n=247). **(B)** Cell clustering and cell types based on patch correlations visualized on a UMAP. Marker pair-wise correlations between patches of the same cell were used as features for UMAP and clustering of HUC, HBM, and HCH. **(C)** Cell clustering and cell types based on network variability visualized on a UMAP. Mean and standard deviation of connection between marker pairs were used as features for UMAP and clustering of HUC, HBM, and HCH. **(D)** Comparison of separability of RNA count, patch correlation, and neighborhood network variability. Fowlkes Mallows score and Rand index were used to quantify the separability of distinct cell populations. Clustering by patch-level correlation and network variability outperforms clustering by single-cell expression when distinguishing the HUC, HBM, and HCH cell types.

### Comparison of cell type separability

With each cell presenting a unique patch correlation, and each patch presenting a unique spatial neighborhood network, we utilized Uniform Manifold Approximation and Projection (UMAP) plots to visualize and cluster cells (**Fig. 8**). First, the cells were clustered based on single-cell RNA count (**Fig. 8A, S11**). The counts were generated by segmenting each cell and counting the detected RNA dots within the segmentation mask. The single-cell RNA count generates a single-cell transcriptomics profile similar to single-cell RNA sequencing (**Fig. S12**). Then, the cells were clustered based on subcellular patch expression correlation (**Fig. 8B**). Lastly, the cells were clustered by the variability of subcellular spatial neighborhood networks as measured by the mean and variance of connectivity between each pair of markers (**Fig. 8C)**. The UMAPs highlight that all three sets of features can separate HBM, HUC, and HCH. Further comparison using Fowlkes Mallows score and Rand index indicates that both patch correlation and network variability separate HBM, HUC, and HCH better than single-cell RNA expression, indicating the potential to discover additional MSC subpopulations masked by single-cell gene expression **(Fig. 8D)**.

### Subcellular structure and RNA spatial neighborhood network

RNA localization is associated with proteins and organelles.^62, 65^ Localization of *COL1A1* on the endoplasmic reticulum (ER) has been found to regulate the expression of *COL1A1.*^65^ We examined the localization of *COL1A1* in ER by applying spaGNN analysis to RNA-FISH and immunofluorescence (IF) combined staining. *ConA* staining enables visualization of the ER and *WGA* works similarly for the Golgi apparatus. We applied the same subcellular RNA density patch and spatial neighborhood network analysis to study the spatial relationships of RNA and protein (**Fig. 9A**).

**Figure 9.**
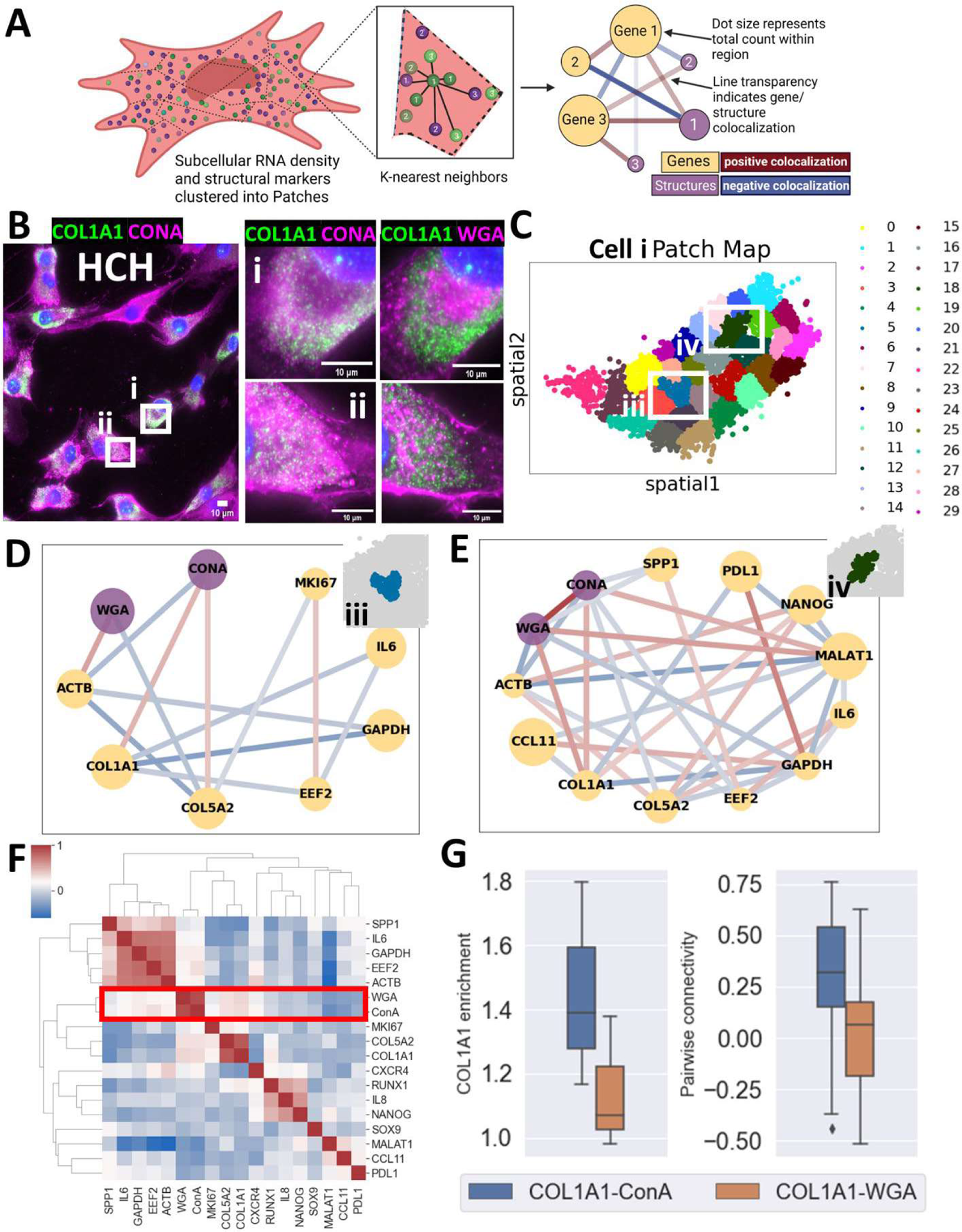
RNA-cellular structure neighborhood network detects *COL1A1* colocalization with ER-targeting *ConA* staining. **(A)** Application of spaGNN to detect RNA-cellular structure colocalization. Subcellular patches were detected using Leiden clustering algorithm. Mean intensity of RNA image around the transcripts indicates the local enrichment of the marker. Correlation between RNA and protein were studied at both patch and local neighborhood level. Correlation at local neighborhood level was visualized as neighborhood networks. **(B)** *COL1A1* overlaid on *ConA* and *WGA* images in 2 regions of HCH highlighted as i and ii. Image shows *COL1A1* localization on *ConA* positive regions and *WGA* negative regions. **(C)** 30 patches of cell highlighted in **Bi**. Patch correlation and neighborhood network were analyzed according to the subcellular patches shown. **(D)** RNA-protein neighborhood network of a patch near the nucleus. *COL1A1* shows positive correlation with *ConA*, and *ACTb* indicates positive correlation with *WGA*. **(E)** RNA-protein neighborhood network of a patch in the cytosolic region. *COL1A1* is suppressed compared to the patch in **D**. The patch shows positive pair-wise correlation between *COL1A1*, *ConA*, and *WGA*. **(F)** Patch correlation of cell highlighted in **Bi**. Correlation between protein markers and RNA are highlighted. **(G)** Comparison of *COL1A1*-*ConA* and *COL1A1*-*WGA* colocalization. The differential colocalization was first analyzed by comparing *COL1A1* enrichment on *ConA* and *WGA* positive regions (Left). *COL1A1* shows higher enrichment in *ConA* positive regions than *WGA* positive regions. The difference in colocalization was confirmed by comparing average connectivity in neighborhood graphs of *COL1A1*-*ConA* and *COL1A1*-*WGA* (Right).

We observed that *COL1A1* tends to colocalize with *ConA*, and inversely colocalize with *WGA* in HCH (**Fig. 9B**). We then expanded the spaGNN analysis to examine the localization of *COL1A1* relative to *ConA* and *WGA*. Structural marker enrichment around an RNA is needed to adapt the spaGNN analysis. A transcript’s associated region is defined as the 3×3 pixels around the transcript. The sum of the pixel intensity of the transcript’s associated region indicates the enrichment of the marker around the transcript. Subcellular patches are defined in the same fashion as previous analyses (**Fig. 9C**). Average marker enrichment around all transcripts in the same patch is defined as the enrichment of the marker of the patch. Within the same patch, 10-nearest neighbor analysis were conducted to find the connectivity between genes and structural markers with the average enrichment of marker around all transcripts indicating marker enrichment levels in the local neighborhood. *ConA* and *WGA* show region-specific correlation at the sub cellular patch level (**Fig. 9D, 9E**). In a *COL1A1* enriched patch, *COL1A1* was highly colocalized with *ConA,* while *β-actin* was highly colocalized with *WGA* (**Fig. 9D**). In a *COL1A1* sparse patch, *COL1A1* was highly colocalized with *WGA* and *ConA* (**Fig. 9E**). Patch correlation among all patches of the same cell were also computed, yet *COL1A1* and *ConA* correlation was not detected (**Fig. 9F**).

We then examined two different criteria among multiple cells to generalize the finding. The first is *COL1A1* enrichment in *ConA* and *WGA* positive regions (**Fig. 9G left**). *ConA* and *WGA* images were thresholded manually, and enrichment of *COL1A1* in marker-positive regions is shown. Further, the average *COL1A1*-*ConA* and *COL1A1*-*WGA* connectivity among all patches of cell i and cell ii (**Fig. 9B**) were compared (**Fig. 9G right**). Both plots show increased colocalization between *COL1A1* and *ConA* than *COL1A1* and *WGA*.

### Edge localization of cytokine gene expression in MSCs

The surface topography of bioreactors used for MSC expansion modulates cytokine secretion profiles of individual stem cells^66^. Bioreactor-grown MSCs exhibit unique morphologies and correlate with a range of anti and pro-inflammatory cytokines. Additionally, the cell density of cultures modifies gene expression of single MSCs due to inter-cellular signaling and communication.^67^ Efficient, spatially resolved secreted factors gene expression can be identified using multiplexed gene expression analysis maps. Such spatial cytokine maps will assist in identifying inflammatory effects of stem cell transplantation and tissue regeneration therapies.

Cytokine release mechanisms are coordinated by intricate spatio-temporal mechanisms guided by cell-to-cell interactions and transient secretion dynamics.^68^ Immunostaining of cytokine proteins yields a portion of pre-release and/or short-term storage cytokine molecules in cells.^69^ Thus, secretion profiles are commonly performed in culture supernatant using multiplexed antibody arrays and Luminex assays, masking the heterogeneity across MSCs.^70, 71^ We reason that gene expression measurements of cytokines will be an important, potentially superior metric to define MSC secretory potential when compared to traditional direct protein measurements.

We applied the spaGNN pipeline to cytokine genes in MSCs with a modified patch definition to extract distinguishing features of cells with edge-localized cytokine RNA (**Fig. 10A**). Three cytokine genes, *IL8*, *IL6*, and *CCL11*, were measured. In some HUCs, cytokine RNAs, especially *IL8* and *CCL11*, are located along the cell membrane (**Fig. 10B, S13**). Contour patches were parallel to the cell membrane with a width. The local neighborhood was reduced to the transcript and its four nearest neighbors with a maximum distance of 5μm (**Fig. 10C**). The maximum distance was set to eliminate transcripts with large distances in between forming a neighborhood.

**Figure 10.**
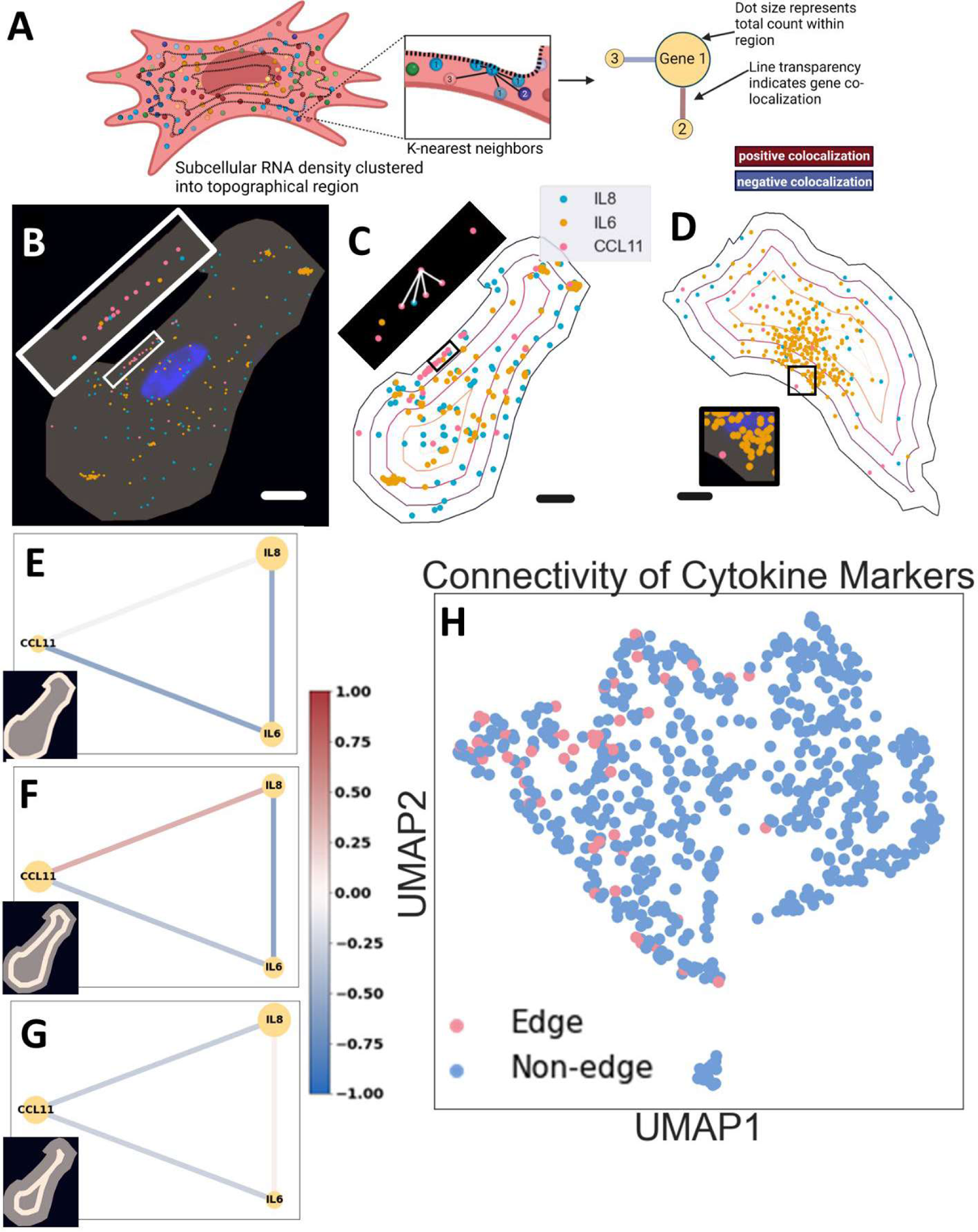
Contour patch model resolves edge-localized gene neighborhood networks. **(A)** Modified spaGNN pipeline to analyze cytokine RNA localization. Contour patches were defined by distance from edge of the cell. The same neighborhood network analysis was applied to each patch. **(B)** Edge localization of *CCL11* transcripts along cell edge. The scatter plot overlaid on cell mask indicates subcellular localization of detected RNA transcripts. Contour patch representation of single cells. RNA transcripts were assigned to contour patches based on their distance to the edge of the cell. Callout contains Nearest neighbors of an RNA transcript in the patch closest to the cell edge, highlighted by the black box. Only the center RNA and 4 nearest neighbors were included in the local neighborhood. Further, only transcripts within 5μm can be considered as neighbors of the central RNA. **(C)** Contour patch model of a cell without edge-localization of cytokine transcripts. **(D)** The neighborhood network of outermost contour patch. The cell shows *CCL11* edge-localization pattern. *IL6* shows negative correlations with both *IL8* and *CCL11*. **(E)** The neighborhood network of contour patch 1 region from cell edge. *IL8* shows moderate positive correlation with *CCL11*. *IL6* shows negative correlations with *IL8 and CCL11*. **(F)** The neighborhood network of patch 2 regions from cell edge. *IL8* shows moderate positive correlation with *IL6*. *CCL11* shows negative correlation with *IL8* and *IL6*. **(G)** UMAP of marker pair-wise connectivity of contour patches used as features. Cells with and without edge localizations of cytokine genes are highlighted in red and blue, respectively.

In a cell showing *CCL11* transcript concentration along the cell membrane and a cell lacking edge RNA localization feature, an equal distance plot shows the patches and RNA localization (**Fig. 10C, 10D**). In the outermost patch of the cell, *IL6* shows a negative correlation with *IL8* and *CCL11* (**Fig. 10E**). *IL6-CCL11* negative correlation is conserved through all patches of the cell, however *IL8-CCL11* and *IL8-IL6* correlation vary (**Fig. 10E, 10F, 10G, S14)**. Cells with both *IL8* and *CCL11* edge localization show different neighborhood networks (**Fig. S15**). Cells were visualized in a UMAP based on the connectivity of cytokine markers in the outermost patch (**Fig. 10H)**. Cells with edge localization of cytokine RNA were also highlighted in gene expression, RNA density patch correlation, and neighborhood network variability UMAPs (**Fig. S16**).

## Discussion

The presented spaGNN workflow identifies cellular and subcellular heterogeneity *in vitro*. However, the biological mechanisms that govern such subcellular RNA organization require further investigations. We reason that several factors may lead to unique subcellular distributions of RNA. Protein secretion and distribution may be influenced by the relative position of the translation unit to one of many possible interacting features. For example, *COL1A1* is secreted and deposited into the extracellular matrix; if *COL1A1* RNA were colocalized with the ER, utilizing the attached translation enrichment units and subsequent Golgi apparatus delivery may increase secretion efficiency^26, 54^. Colocalization of RNA could also potentially be attributed to RNA-structural protein binding.^26^ These RNA-protein interactions can serve both structural purposes and post-transcriptional regulation purposes, among others.^72^ The localization of RNA may also assist the function of the translational product; for example, *β-actin* mRNA can localize near adhesion points to assist cell migration and adhesion.^31, 73^ Similar effects of localized mRNA have been observed in neurons to maintain neural plasticity, a crucial factor in learning and memory.^74^ We also speculate that specific localization of mRNA may increase energy efficiency. By transporting mRNA to various subcellular locations to translate proteins where they are needed, cells avoid the expensive transport of large, complicated molecules in favor of much smaller mRNA structures.

Additionally, clonal heterogeneity has not been assessed in these samples, making it difficult to decipher the genetic purity of MSC colonies. Multiplexed RNA imaging methods can be applied to MSCs after engineering each stem cell using a barcoded library of lentiviral vectors.^75^ Simple cell trackers can also be used to identify colony-forming units of individual MSCs, but these methods cannot be effectively utilized in the study of clonal mixing and evolution.^76^ Alternatively, single-cell time-lapse imaging can map the differentiation process by phase-contrast microscopy followed by multiplexed, spatial gene expression measurements.^77^ Synthetic lineage recording methods can also be applied to decipher clonal heterogeneity, but these molecular barcoding methods are difficult to execute in human stem cells.^78^ Other molecular analysis methods such as ATAC-seq can also be incorporated into the clonal mapping of single MSCs.^79^

While this MSC study was used to profile and quantify only HBM, HUC, and HCH from one donor per cell type, this image-based screening platform may be fully automated to enable a high-throughput screening assay for comparative analysis of many donors and tissue-specific MSCs. Such an MSC monitoring assay could prove essential in biomanufacturing processes such as end-point measurements and donor screenings, providing a potential critical quality attribute (CQA) to standardize functional MSC products. Integrating the aforementioned analytical methods in addition to the spaGNN pipeline would enable discovery of biological mechanisms in MSCs not observable with bulk transcriptomics.

Other stem cell sources such as induced Pluripotent Stem Cells (iPSCs) may also benefit from the spaGNN framework as a method to identify cell states, differentiation potential, and immunomodulatory functions. Complementary RNA-seq assays may also be used to predict the identity and function of individual stem cells. Data integration can be performed using developing multimodal data integration techniques^80, 81^. In short, the spaGNN image-based spatial omics framework assists in the identification of stem cell subpopulations utilized in cell-based regenerative treatments as well as the broader study of stem cell biology.

### Summary

The SpaGNN workflow sheds light on cellular and subcellular organization of single HBMs and HUCs. The presented computational clustering methods reveal unique subcellular distributions of RNA and structural features. The pattern of RNA spatial distribution is modeled by a transcript spatial enrichment analysis, while subcellular RNA density patch correlation indicates RNA colocalization at the subcellular region level. Machine learning algorithms detect significant differences in patch-level correlation between cell types, and local nearest neighbor analysis reveals heterogeneous spatial co-expression of RNA across patches in the same cell. Single-cell RNA counts, pair-wise patch-level correlation, and variability of local neighborhood network distinguish between HBM, HUC, and HCH cells, with patch-level correlation and network variability providing stronger separability. By incorporating structural marker enrichment in each patch, the spaGNN pipeline detects RNA-structure colocalization relationships. Further application of spaGNN into the analysis of cytokine genes demonstrates cellular and subcellular heterogeneity of cytokine gene localization. The spaGNN framework is an emerging single cell spatial omics tool for characterizing stem cells and informing biomanufacturing pipelines.

## STAR Method

### Cell culture

Bone marrow-derived MSCs (BM-MSCs) and umbilical cord-derived MSCs (UC-MSCs) were purchased from RoosterBio, Inc. (**Table S1**). The culture media consisted of 89% L-glutamine supplemented α-MEM media (Cat # 12561-049), 10% heat-inactivated fetal bovine serum (HI-FBS), and 1% penicillin-streptomycin (Cat # P4333). The culture media was first mixed, then filtered before use. HBM and HUC were cultured in T-75 flasks with 15mL of culture media. Cell passages were performed when cells reach 75% confluency using 5mL of Trypsin LE cell detachment media (Cat # 12605-010) per T-75 flask for 5 minutes. After confirming cell detachment, 5mL of media was added to each flask to neutralize the trypsin. Cell suspension was then collected and centrifuged at 280g for 6 minutes. Cells were then resuspended in their respective culture media. Suspended cells were then seeded on collagen-coated glass coverslips. The cells were then cultured on coverslips for 24 hours before fixation.

Cryopreserved human primary chondrocytes (HCH) (C-12710) and chondrocyte culture media (C-27101) were purchased from PromoCell, GmbH. The media consisted of basal media and supplement mix. The basal media and supplement mix were mixed at a ratio specified by the instruction manual. Cell passages were performed when cells reach 75% confluency using Trypsin LE cell detachment media (Cat # 12605-010) at room temperature and resuspension in respective culture media after centrifugation at 220g for 3 minutes. Suspended cells were then seeded on collagen-coated coverslips. The seeded cells were cultured for 24 hours before fixation.

Cells were fixed with 4% paraformaldehyde fixation buffer (4% paraformaldehyde (Cat # 28908) with 1X DPBS (diluted from 10x DPBS (REF D1408) with DNase, RNase free water (Cat # 10977-015) for 10 minutes. The samples were then washed with 1x DPBS (Cat # D8537) twice and stored in cold ethanol at −20°C overnight for permeabilization.

### Multiplexed RNA detection

The HCR staining follows the protocol provided by Molecular Instruments RNA-FISH on mammalian cells on slides protocol. After permeabilization, the samples were air-dried for 10 minutes and washed with 2x SSC buffer 3 times. The samples were then incubated in a 300μL pre-warmed HCR probe hybridization buffer (LOT# BPH02023) at 37°C for 30 minutes for prehybridization. 3L of HCR probes for desired targets were then added to 300μL of pre-warmed probe hybridization buffer. The prehybridization buffer was aspirated, and probe dilute was added to the samples. The samples were incubated at 37°C for 12-16 hours. The samples were then washed with 300μL of pre-warmed probe wash buffer (LOT# BPW02922) for 5 minutes at 37°C utes at room temperature at room temperature for 30 minutes for pre-amplification. 6μL of each HCR amplifier hairpins were then snap-cooled down to room temperature in a dark drawer for 30 minutes. The snap-The pre-amplification buffer was then aspirated, and diluted hairpins were added to the sample. The samples were then incubated for 1 hour and 15 minutes. The amplification **s**olution was then aspirated, and the samples were washed with 5x SSCT for 5 minutes at room temperature 5 times (see HCR protocol for cells on slide).

The samples were then mounted in an antifade mounting buffer containing Tris-HCL (20 mM), NaCl (50 mM), glucose (0.8%), saturated Trolox (Sigma, 53188-07-1), pyranose oxidase (Sigma: P4234), and catalase (Sigma, 9001-05-2, 1:1000 dilution).

To remove fluorescent mRNA signals, we used RNase-free DNase I (Sigma, 04716728001). After imaging, the samples were first washed with 2x SSC twice. We then diluted 50μL of 10x concentration incubation buffer in 450μL of R Nase-free water to 1x concentration. The samples were then incubated with a 1x incubation buffer at room temperature for 5 minutes. Then 10 μL of DNase 1, 50 μL of 10x incubation buffer, and 440μL of RNase-free water were mixed. The samples were then incubated in the DNase I mixture for 4 hours at room temperature. After incubation, the samples were then washed with 30% formamide in 2 SSC at room temperature for 5 minutes 3 times. Removal of signals was confirmed under a fluorescent imaging microscope. The samples were then ready to start the next cycle of RNA labeling.

### Image preprocessing

Images of cells in 2D culture all underwent the following preprocessing before further analysis. Each image contains multiple channels and each channel is composed of images collected at various z-levels. The background of each gene-positive channel was first subtracted using a rolling ball method with a radius of 15 pixels. The maximum intensity projection of each channel was then generated. The projected image was the final image used for subcellular clustering and counting analysis. For the DAPI channel, the maximum intensity projection was generated from the raw image. Previous steps were conducted in a script implemented in FIJI, enabling automated high throughput processing.

To apply clustering analysis, the images across multiple cycles were registered using phase cross correlation method in scikit-image^82^. After detecting the translational and rotational shifts of the images, the reversed shifts were then applied to images, and overlapping regions were then cropped and saved.

RNA HCR-FISH signal was detected by finding local maximum in each image above a manually set threshold. The detection algorithm was implemented using local_peak_max() function in the scikit-image package. Parameters adjusted in local_peak_max() were absolute threshold and minimum distance between local peaks. The parameters were adjusted by each gene. The custom-developed code detected all peaks in the image, then applied the segmentation mask to find RNA counts and locations per cell.

### Subcellular clustering

Pixels within cells were first selected from the images according to a manually segmented cell mask. Selected pixels were down-sampled by 20 folds to form a represented sub-sample of the whole dataset. The selected pixels were normalized to mean of 0 and standard deviation of 1. Then, a Leiden clustering algorithm was applied to the down-sampled dataset to assign a cluster label to each pixel. Finally, pixels of the entire dataset were assigned to the cluster of most likely neighbors among 30 nearest neighbors in pixel feature space in the down-sampled dataset.

In K-means clustering, the extracted pixels were first normalized to mean of 0 and standard deviation of 1. K-means clustering was then applied with K values determined by finding “elbow” points on the within-cluster sum square error versus K-value plot. Pixel distributions for each cluster were displayed with heat maps.

### Subcellular RNA distribution statistics

The center of a cell is defined by the center of mass of the cell mask. The center of mass of the mask is the average location of all pixels in the mask, and was calculated by the following equation:

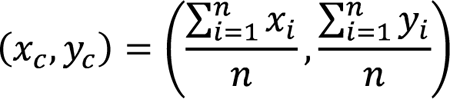

where *x_i_* and *y_i_* are positions of pixels in the segmentation mask.

The distance between gene and center of mask is the Euclidian distance, and was calculated by the following equation:

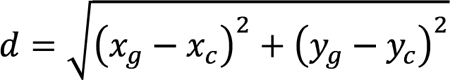

where *x_g_, y_g_*, are the position of genes.

Enrichment of RNA at distances from center of the cell was then calculated. The subcellular distribution of RNA was compared between cells using Kolmogorov-Smirnov test implemented in scipy.stats package.

### Subcellular patch analysis

Positions of all detected RNA transcripts were combined and clustered using the Leiden clustering algorithm with a resolution factor of 1. The Leiden clustering algorithm was implemented by scanpy.tl.leiden() function in Scanpy package^61^. RNA transcripts of the same cluster were considered in the same patch. Copy numbers of genes in each patch were then counted. Pearson’s correlation between genes were then computed among patches.

### Subcellular local neighborhood analysis

Local neighborhoods were detected by applying the k-nearest-neighbor algorithm (k-NN) to positions of all RNA in the same patch. We utilized sklearn.neighbors.NearestNeighbors package implemented in scikit-learn^83^. Each neighborhood consisted of the transcript and its 9 nearest neighbors. Copy numbers of genes in each local neighborhood were counted. Pearson’s correlation between genes were computed among all local neighborhoods in the same patch. The gene neighborhood network is visualized using the networkx package^84^.

### Clustering and visualization of cell spatial transcriptomic profile

In gene expression clustering analysis, the counts of genes were calculated by counting all detected RNA in the same cell mask. In patch correlation clustering, pairwise correlations among patches of each cell were collected. The correlations of different genes were used as features for clustering. In network variability clustering, the mean and standard deviation of connectivity between each pair of genes of the same cell were used as features for clustering.

### RNA-cellular structure colocalization

The enrichment of cellular structure marker around each RNA was extracted by computing the total intensity of 3×3 pixels around the location of the RNA. Mean enrichment around all RNA in the same patch or local neighborhood were computed to indicate marker enrichment of the patch or local neighborhood. Correlations were computed similarly to all previous correlation analysis.

Enrichment measurement in **Fig. 9G** was calculated as follows. *ConA* and *WGA* images were manually thresholded, and divided into marker positive regions and marker negative regions. The proportion of marker positive region in the cell is calculated by the following equation:

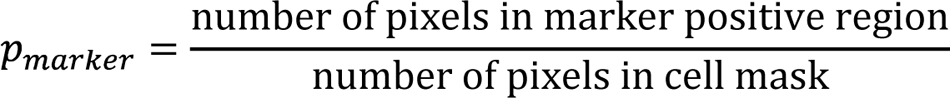

Then, the proportion of *COL1A1* transcript detected in the marker positive regions were calculated by the following equation:

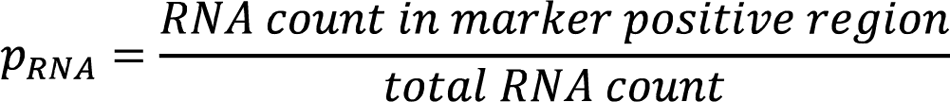

The enrichment of RNA in protein-positive region was calculated by the following equation:

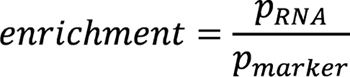

### Cytokine gene spatial distribution

After a cell was segmented, a map of distances from all points inside the cell to the edge of the cell was generated. Distances to the edge of the cell were calculated for each RNA. Cells were then divided into contour patches of 5μm width according to the distance to the edge of the cell. The same local neighborhood correlation analysis as before was conducted for each patch. Local neighborhood was reduced to the transcript and 4 nearest neighbors. Transcripts with distance to the center transcript greater 5μm from the neighborhood. To visualize the distribution of cells with edge localized cytokine RNA, Pearson’s correlation between all local neighborhoods of the outer-most patch were used as features to generate UMAP.

## Acknowledgments

A.F.C. holds a Career Award at the Scientific Interface from Burroughs Wellcome Fund and National Institute of Health K25 Career Development Award (K25AI140783). A.F.C. was supported by start-up funds from the Georgia Institute of Technology and Emory University. This material is based upon work supported by the National Science Foundation under Grant Number. EEC-1648035. A.J.F. is supported by National Science Foundation Center for Cell Manufacturing Technologies Research Experience for Undergraduates (REU) program.

## Author Contribution

Conceptualization: Z.F., A.M., A.C.; methodology, Z.F., A.C.; software, Z.F., T.H., N.Z.; investigation, Z.F.; supervision, A.C.; writing – original draft, Z.F., A.F., T.H., N.Z., A.C.; writing – review & editing, Z.F., A.F., T.H., N.Z., A.C.; visualization, Z.F., A.F, T.H., N.Z.; and funding acquisition: A.C.

## Declaration of Interest

The authors declare no competing interests.

## Data and Code Availability

Example data and custom codes are available at: https://github.com/coskunlab/spaGNN

## Notes

### Competing Interest Statement

The authors have declared no competing interest.

