## Supplemental Material for "Subcellular spatially resolved gene neighborhood networks in single cells"

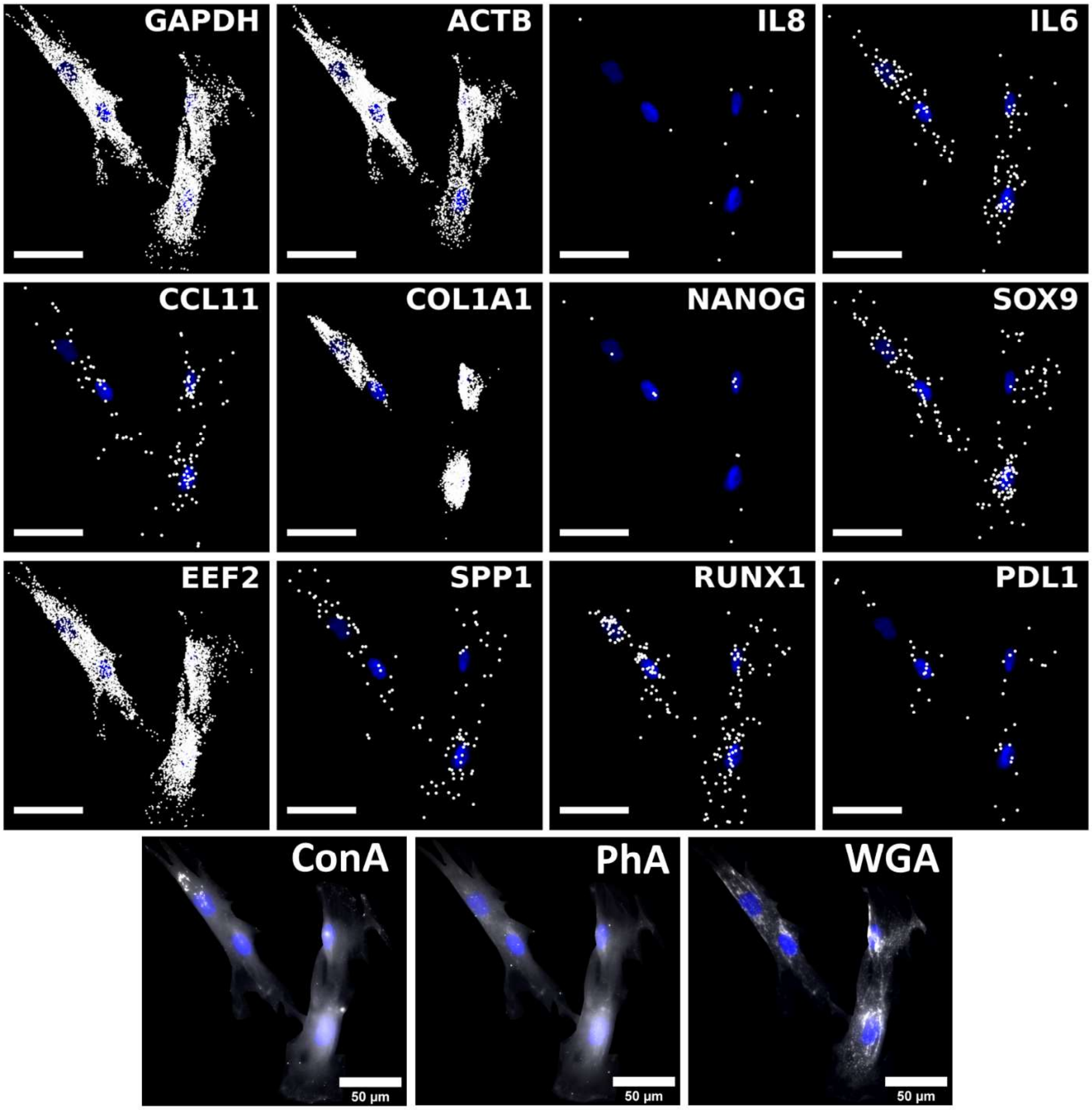

GAPDH
ACTB
IL8
IL6
CCL11
COL1A1
NANOG
SOX9
EEF2
SPP1
RUNX1
PDL1
ConA
PhA
WGA
50 µm
50 µm
50 µm

**Figure S1. Multiplexed HCR generates spatial transcriptomics profile of HBM** Scatter plot showing localization of RNA and adjusted images of protein markers in HBM. The positions of RNA molecules were plotted on the *DAPI* image. *GAPDH*, *ACTB*, *EEF2*, and *COL1A1* are highly expressed, with *COL1A1* showing significant enrichment around the nucleus. Brightness and contrast adjusted *ConA*, *PHA*, and *WGA* images were overlaid on *DAPI* image. Unlabeled scale bars have length of 50µm.

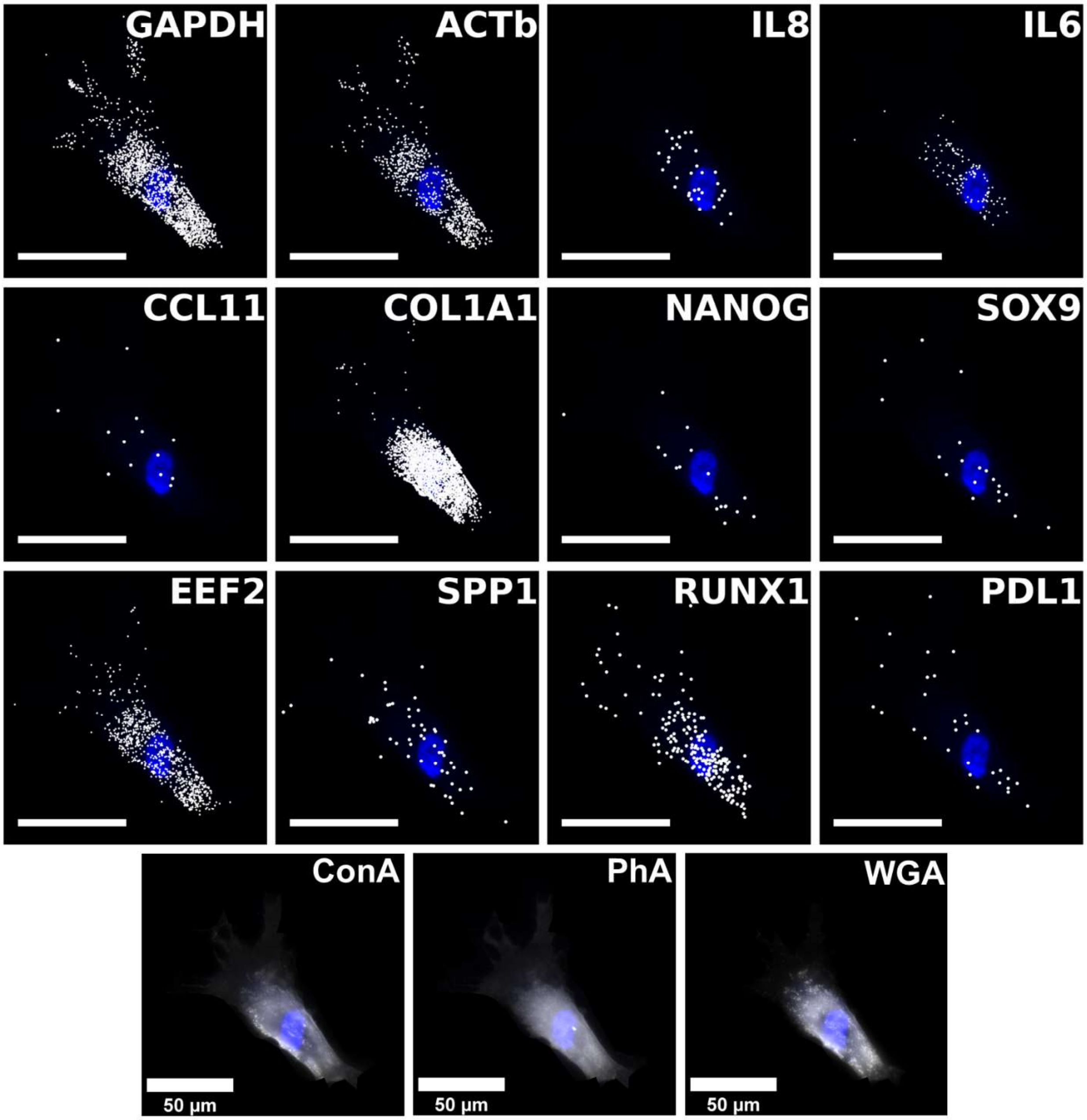
GAPDH
ACTb
IL8
IL6
CCL11
COL1A1
NANOG
SOX9
EEF2
SPP1
RUNX1
PDL1
ConA
PhA
WGA
50 µm
50 µm
50 µm

**Figure S2. Multiplexed HCR generates spatial transcriptomics profile of HUC** Scatter plot showing localization of RNA and adjusted images of protein markers in HUC. The positions of RNA molecules were plotted on the *DAPI* image. *GAPDH*, *ACTB*, *EEF2*, and *COL1A1* are highly expressed, with *COL1A1* showing significant enrichment around the nucleus. Brightness and contrast adjusted *ConA*, *PHA*, and *WGA* images were overlaid on *DAPI* image. Unlabeled scale bars have length of 50μm.

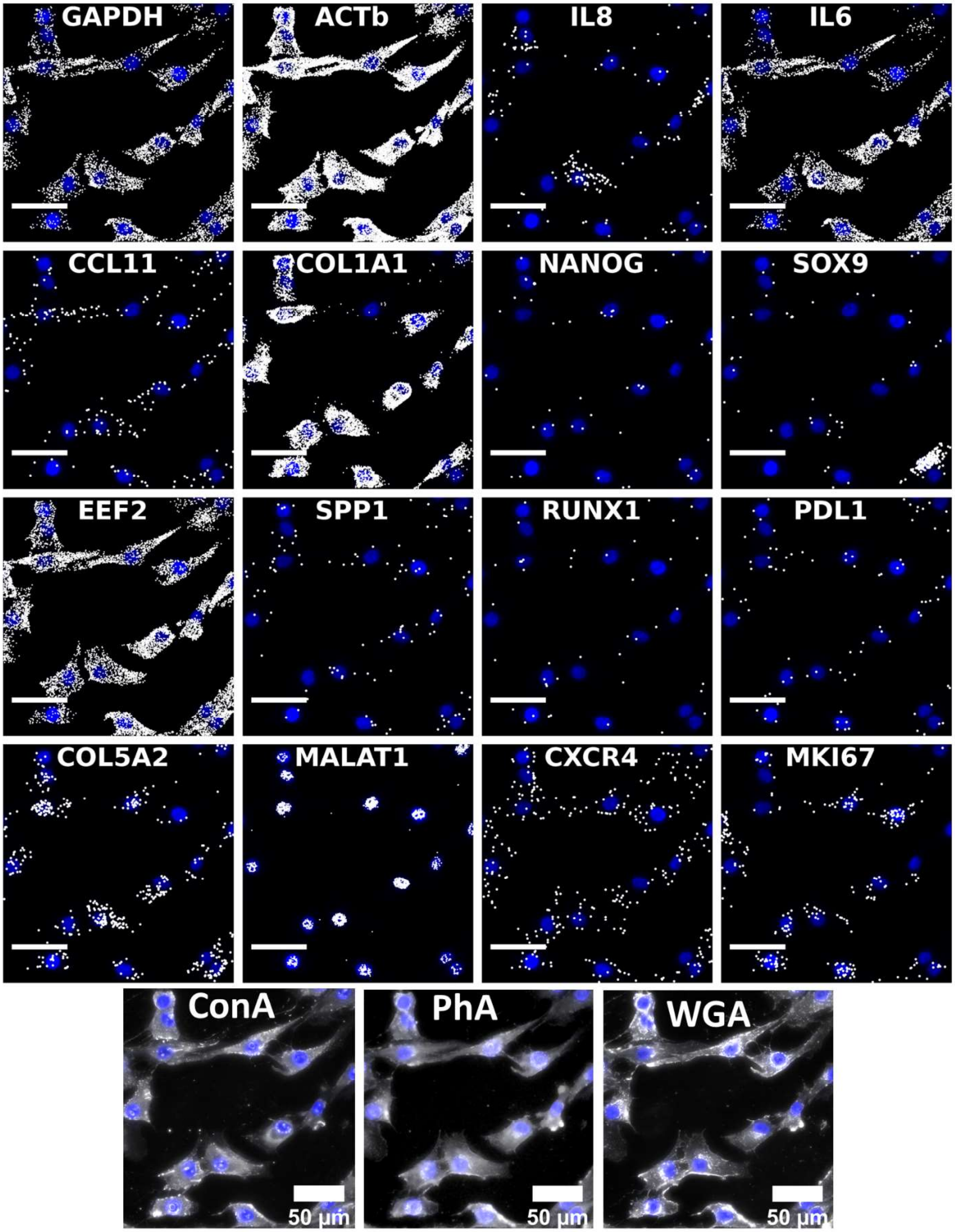
GAPDH
ACTb
IL8
IL6
CCL11
COL1A1
NANOG
SOX9
EEF2
SPP1
RUNX1
PDL1
COL5A2
MALAT1
CXCR4
MKI67
ConA
PhA
WGA
50 μm
50 μm
50 μm

**Figure S3. Multiplexed HCR generates spatial transcriptomics profile of HCH**

Scatter plot showing localization of RNA and adjusted images of protein markers in HCH. Positions of RNA molecules were plotted on the *DAPI* image. *GAPDH*, *ACTb*, *IL6*, *EEF2*, *COL1A1*, and *MALAT1* are highly expressed, with *COL1A1* showing significant enrichment around the nucleus, and *MALAT1* showing exclusive localization in nucleus. Brightness and contrast adjusted *ConA*, *PHA*, and *WGA* images were overlaid on *DAPI* image. Unlabeled scale bars have length of 50µm.

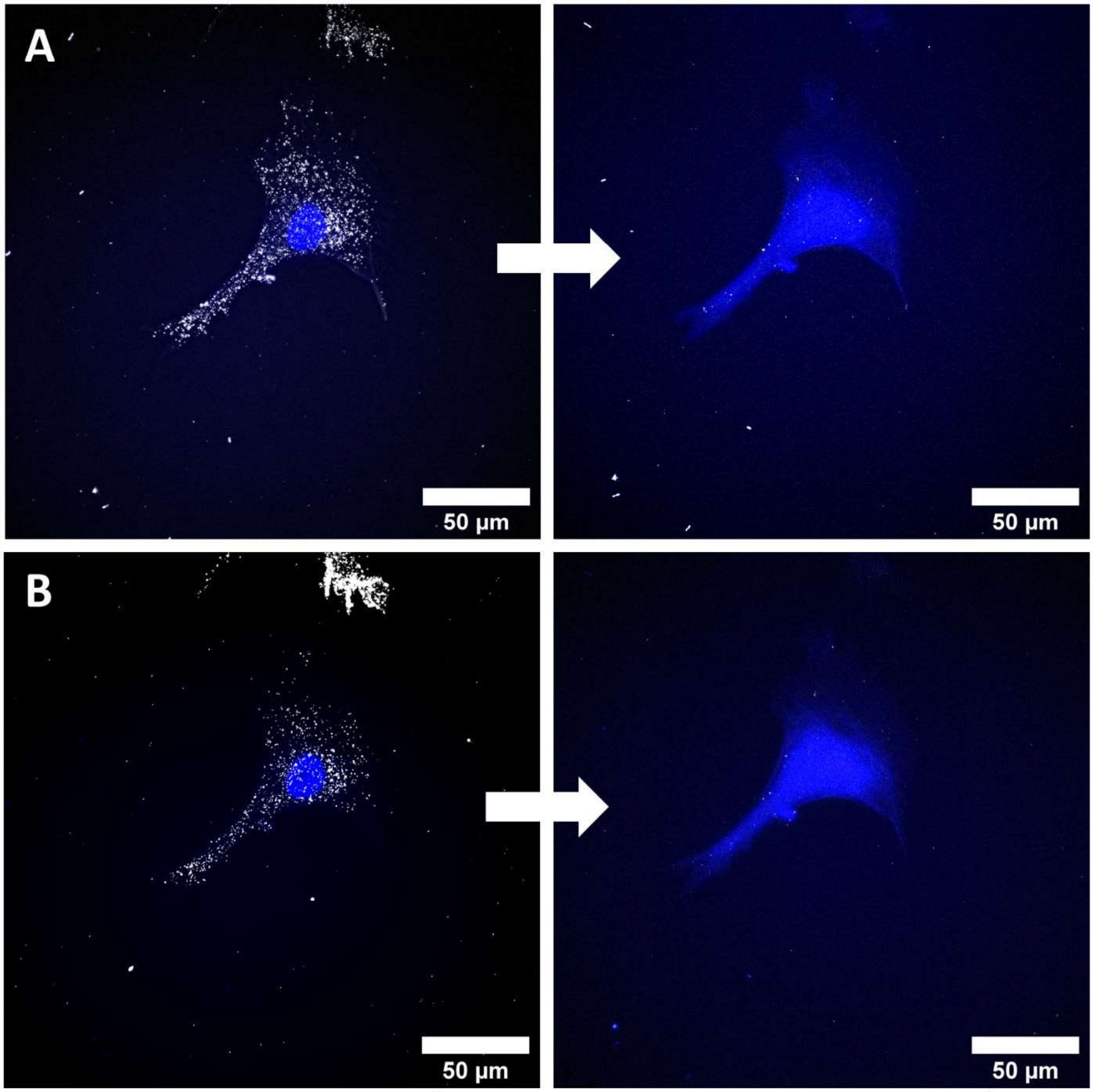


RNA-FISH signal before DNaseI

RNA-FISH signal after DNaseI

**Figure S4. Comparison of RNA-FISH signal before and after DNase I treatment indicates the removal of fluorescent DNA construct**

Removal of HCR constructs fluorescent DNA assembly on RNA. DNase I treatment followed by formamide washes removes fluorescent signal and preserves intact RNA molecules in different channels (A, B).

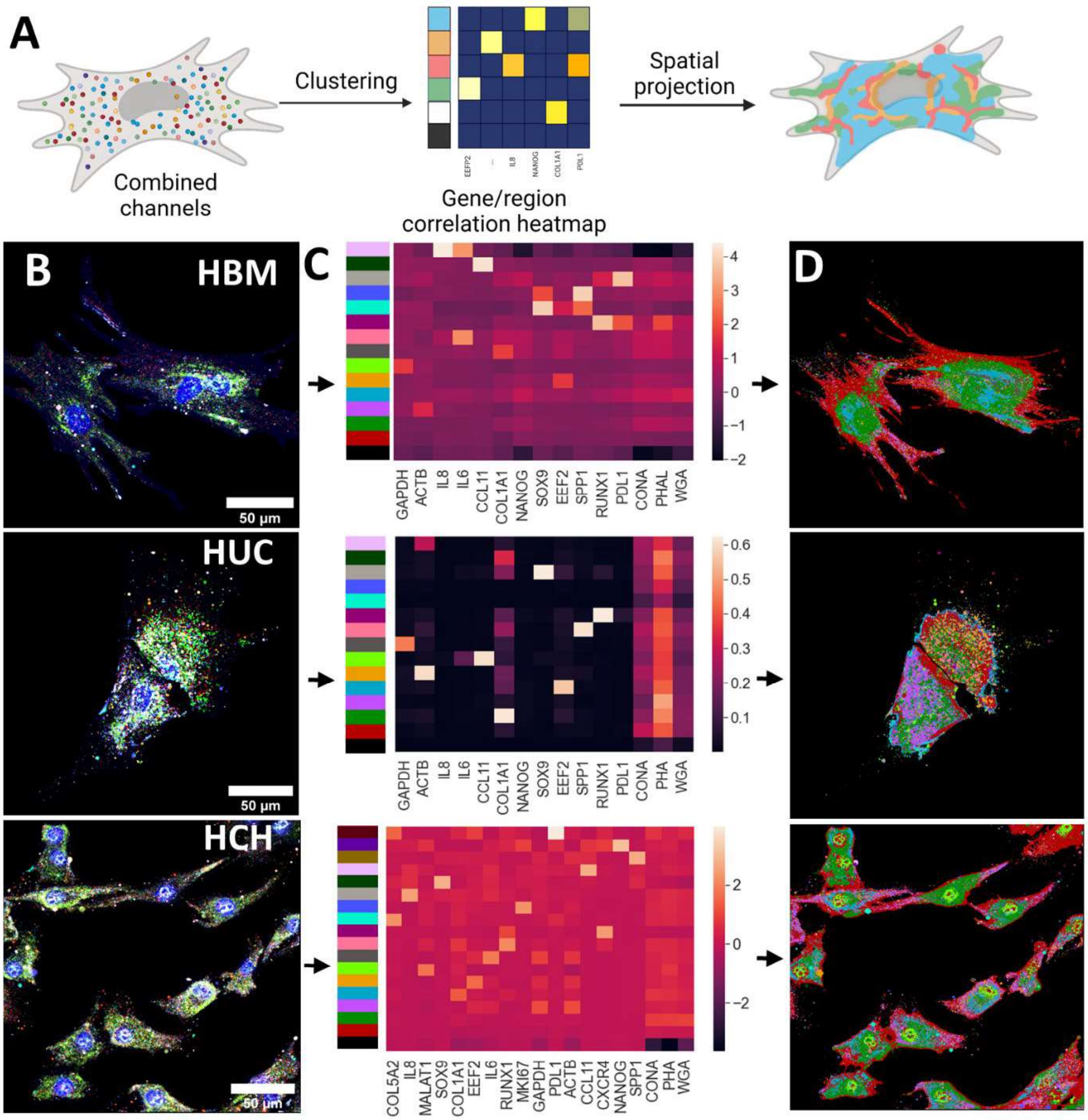
A
Clustering
Spatial projection
Combined channels
Gene/region correlation heatmap
B
HBM
C
D
50 µm
GAPDH
ACTB
IL8
IL6
CCL11
COL1A1
NANOG
SOX9
EEF2
SPP1
RUNX1
PDL1
CONA
PHAL
WGA
HUC
PHA
HCH
COL5A2
MALAT1
MKI67
CXCR4

**Figure S5. Subcellular clustering by single image reveals gradient pattern of gene enrichment region**

**(A)** Subcellular pixel-level clustering of aligned cell images. The image of each cell was analyzed using K-means clustering. The heatmap indicates enrichment of each marker in each cluster. Cells were then pseudo-colored based on the cluster assignment of each pixel.

**(B)** Aligned images of HBM, HUC, and HCH cells.

**(C)** Heatmaps indicating enrichment of markers in each cluster. Pixels from the same image inside cellular regions were collected and clustered using K-means clustering. The cluster centers were defined by computing the mean enrichment of each marker of all pixels assigned to the cluster. The enrichment of markers of each cluster center is shown in the heatmap.

**(D)** Pseudo-colored cell image according to clustering result. Cells were colored based on the cluster assignment of each pixel. Cells present similar gradient patterns of cluster distribution. The same pattern is observed in the same clustering analysis using pixels from multiple images.

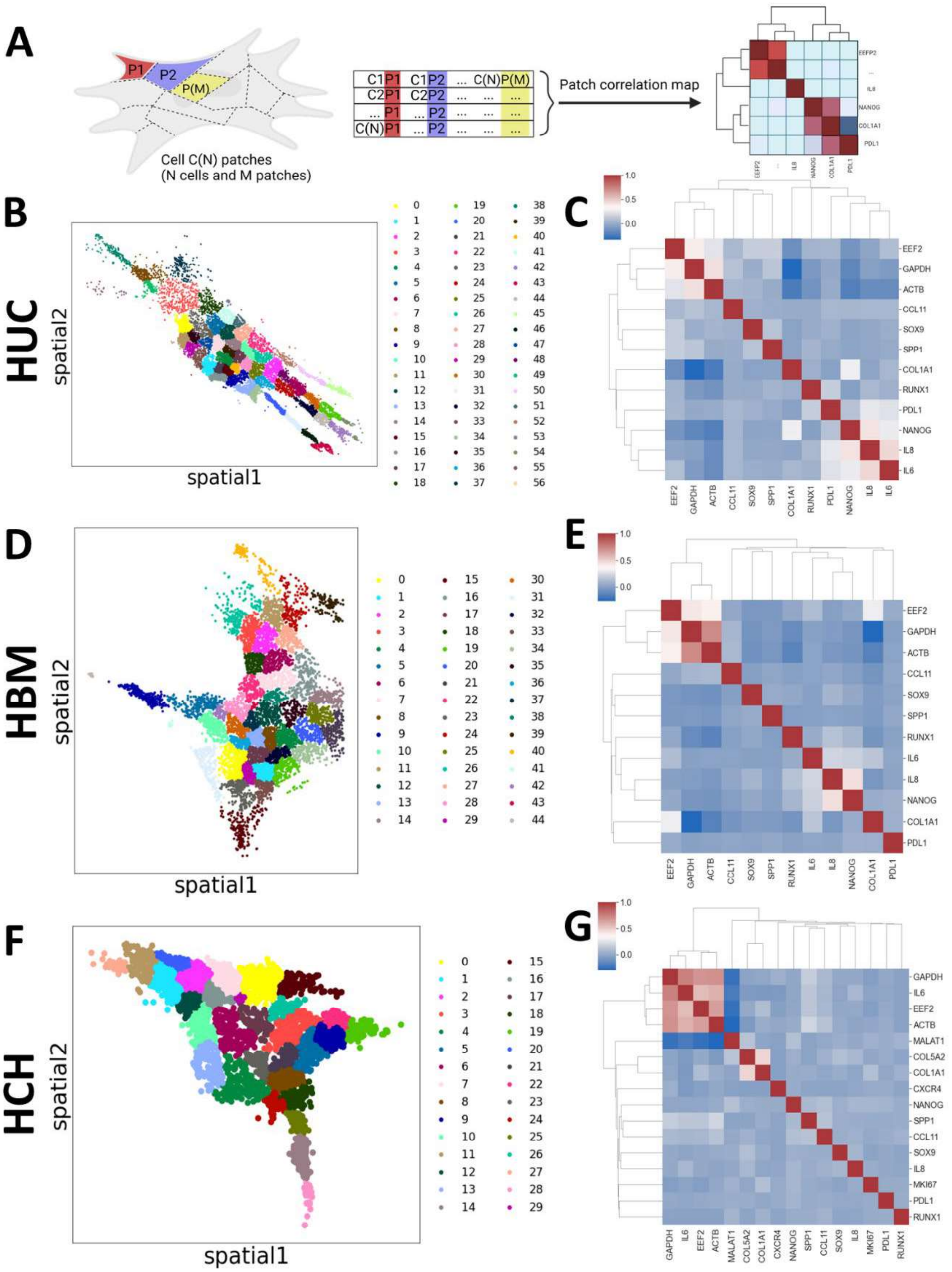
A
P1
P2
P(M)
Cell C(N) patches
(N cells and M patches)
Patch correlation map
B
HUC
spatial2
spatial1
C
EEF2
GAPDH
ACTB
CCL11
SOX9
SPP1
COL1A1
RUNX1
PDL1
NANOG
IL8
IL6
D
HBM
E
F
HCH
G
MALAT1
COL5A2
CXCR4
MKI67

**Figure S6. Subcellular patch correlation of all patches in each cell type indicates differential correlation between cell types**

**(A)** Subcellular pixel-level clustering of aligned cell images. The image of each cell was analyzed using K-means clustering. The heatmap indicates enrichment of each marker in each cluster. Cells were then pseudo-colored based on the cluster assignment of each pixel.

**(B)** Aligned images of HBM, HUC, and HCH cells.

**(C)** Heatmaps indicating enrichment of markers in each cluster. Pixels from the same image inside cellular regions were collected and clustered using K-means clustering. The cluster centers were defined by computing the mean enrichment of each marker of all pixels assigned to the cluster. Enrichment of markers of each cluster center is shown in the heatmap.

**(D)** Pseudo-colored cell image according to clustering result. Cells were colored based on the cluster assignment of each pixel. Cells present similar gradient patterns of cluster distribution. The same pattern is observed in the same clustering analysis using pixels from multiple images.

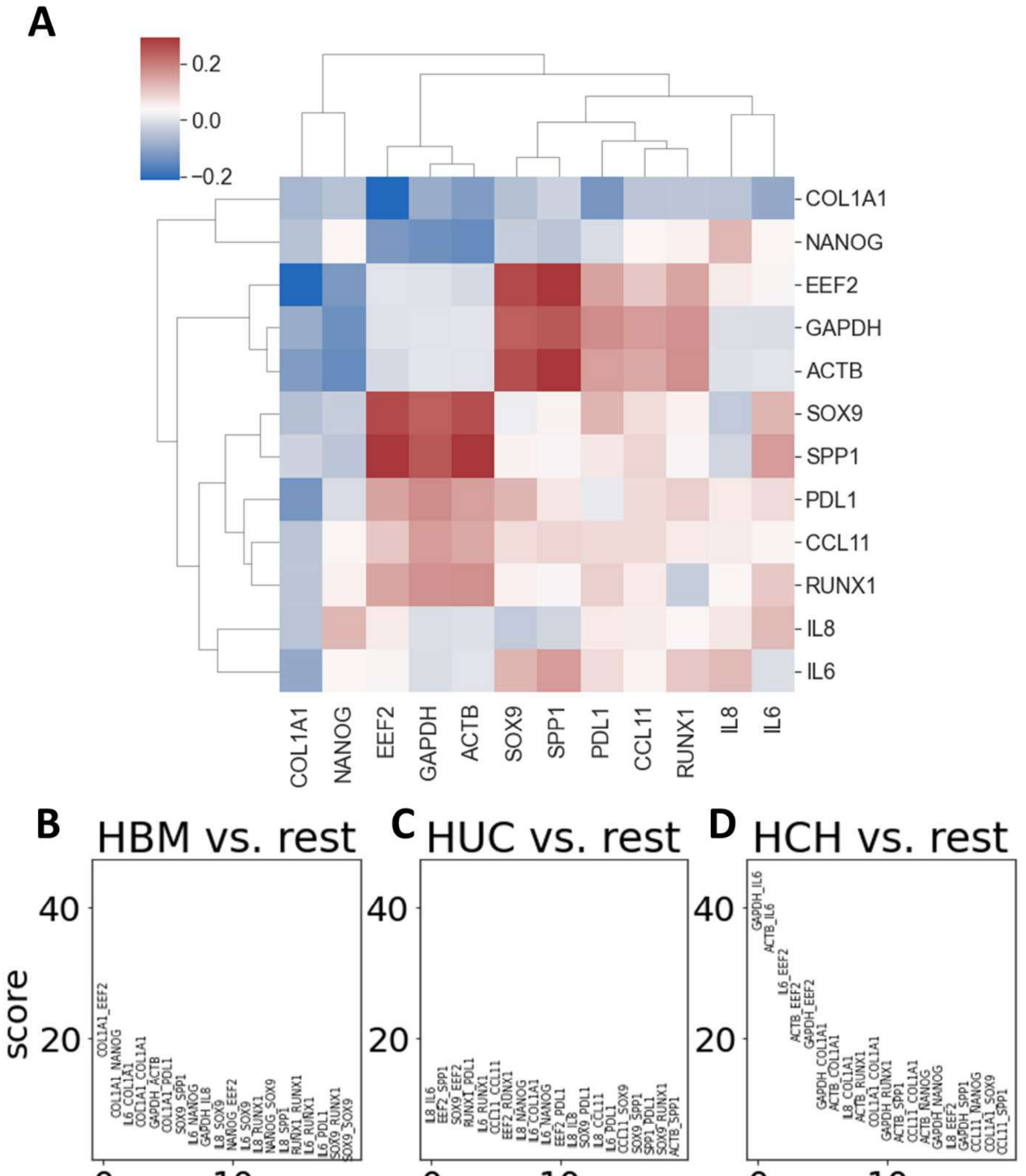
A
0.2
0.0
−0.2
COL1A1
NANOG
EEF2
GAPDH
ACTB
SOX9
SPP1
PDL1
CCL11
RUNX1
IL8
IL6
B
HBM vs. rest
C
HUC vs. rest
D
HCH vs. rest
score
40
20
0
10
ranking
COL1A1_EEF2
COL1A1_NANOG
L6_COL1A1
COL1A1_COL1A1
GAPDH_ACTB
COL1A1_PDL1
SOX9_SPP1
L6_NANOG
GAPDH_IL8
L8_SOX9
NANOG_EEF2
L6_SOX9
L8_RUNX1
NANOG_SOX9
L8_SPP1
RUNX1_RUNX1
L6_RUNX1
L6_PDL1
SOX9_RUNX1
SOX9_SOX9
L8_IL6
EEF2_SPP1
SOX9_EEF2
RUNX1_PDL1
L6_RUNX1
CCL11_CCL11
EEF2_RUNX1
L8_NANOG
L6_COL1A1
L6_NANOG
EEF2_PDL1
L8_IL8
SOX9_PDL1
L8_CCL11
L6_PDL1
CCL11_SOX9
SOX9_SPP1
SPP1_PDL1
SOX9_RUNX1
ACTB_SPP1
GAPDH_IL6
ACTB_IL6
L6_EEF2
ACTB_EEF2
GAPDH_EEF2
GAPDH_COL1A1
ACTB_COL1A1
L8_COL1A1
ACTB_RUNX1
COL1A1_COL1A1
GAPDH_RUNX1
ACTB_SPP1
CCL11_COL1A1
ACTB_NANOG
GAPDH_NANOG
L8_EEF2
GAPDH_SPP1
CCL11_NANOG
COL1A1_SOX9
CCL11_SPP1

**Figure S7. PCA analysis helps identify differences in patch correlation between cell types**

**(A)** Loading of patch-level correlation in PC2. Blue color indicates negative loading of the marker-pair correlation in PC2, and red color indicates positive loading of marker-pair correlation in PC1

**(B)** HBM versus rest separability ranking. High scores indicate the difference between correlation of HBM and the rest.

**(C)** HUC versus rest separability ranking.

**(D)** HCH versus rest separability ranking. Noticeably, most high-ranking correlations involve *IL6*.

**HBM**

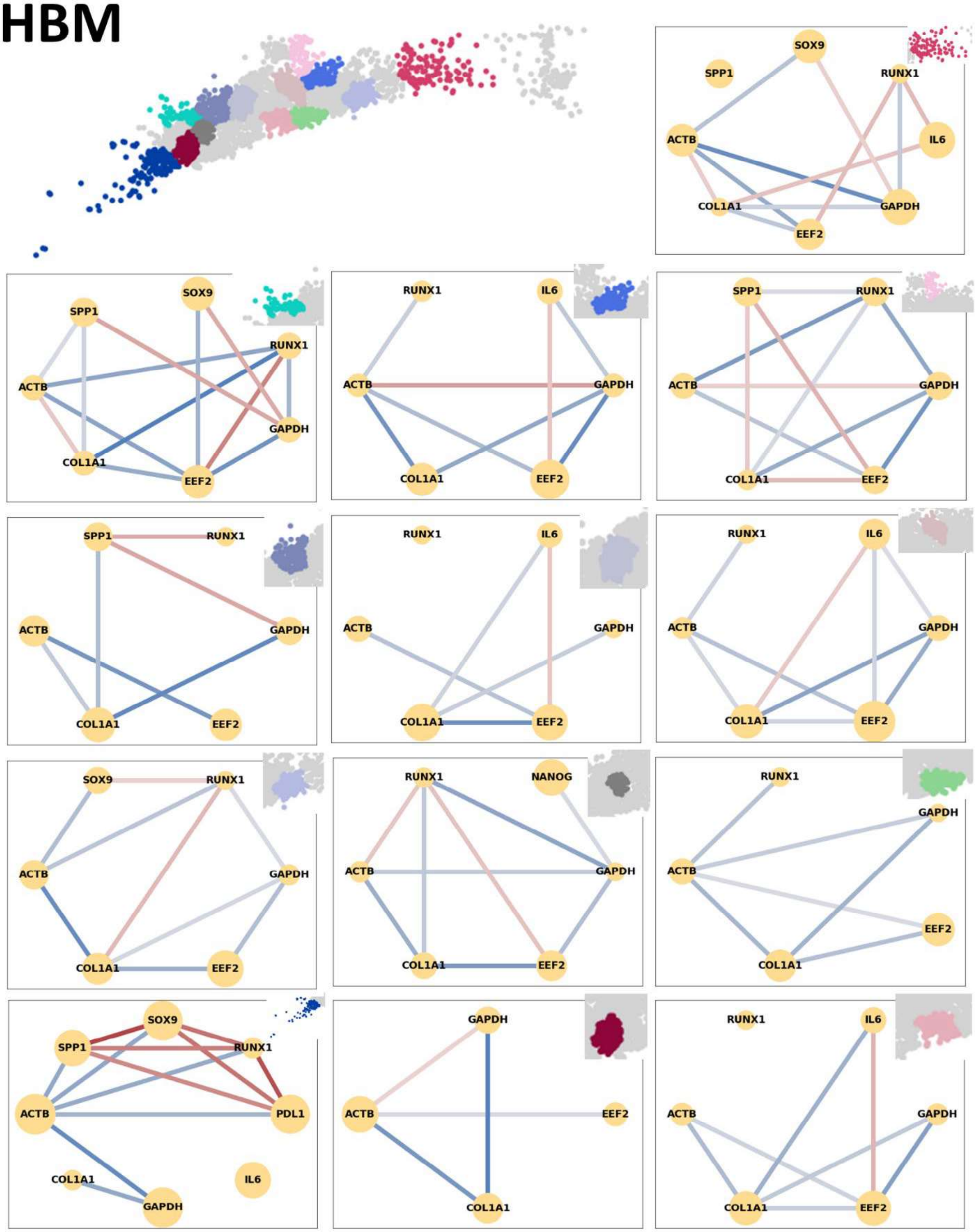

**Figure S8. HBM showing heterogeneous spatial neighborhood networks**

Neighborhood network of a single HBM cell, revealing heterogeneity in different patches.

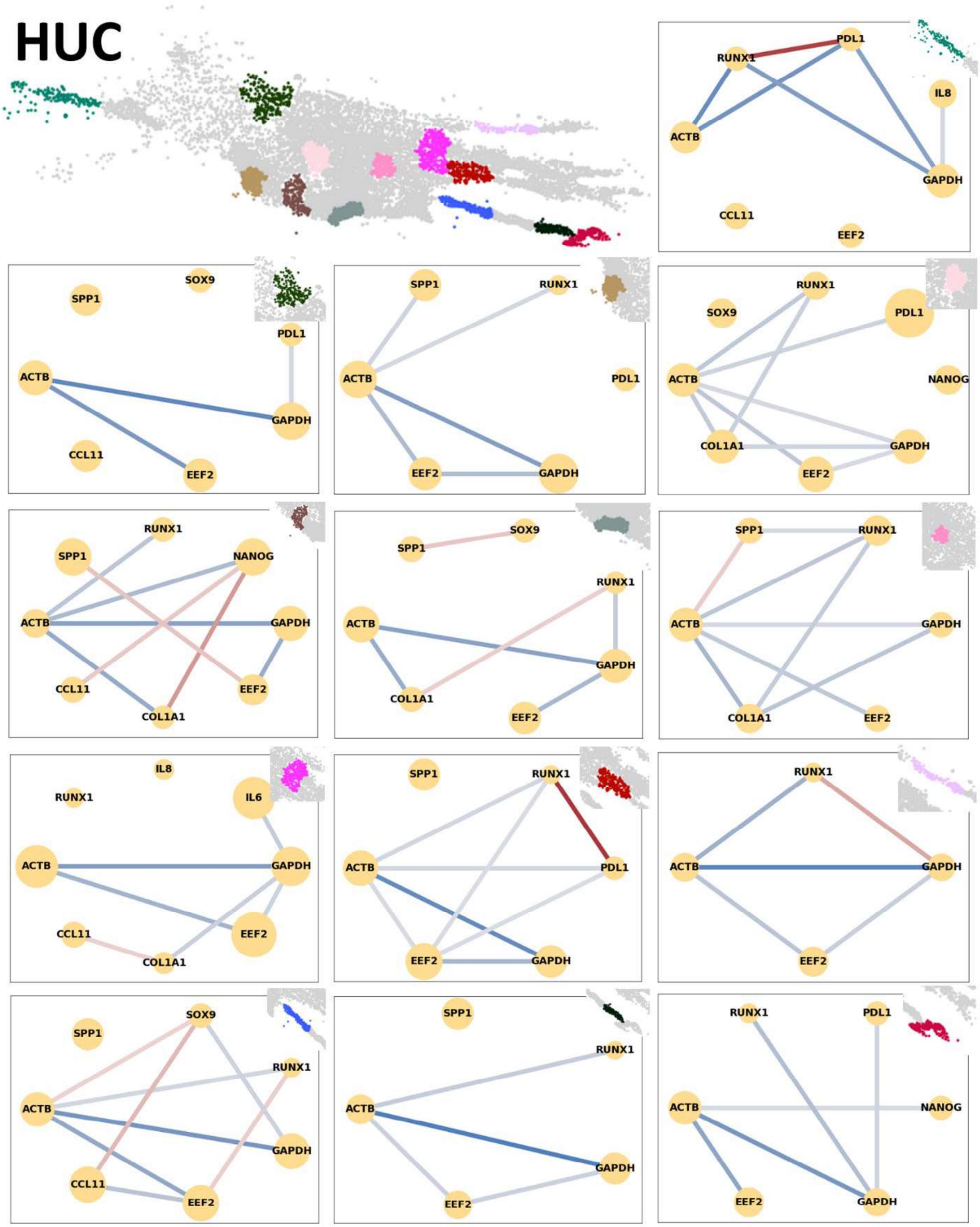
HUC
RUNX1
PDL1
IL8
ACTB
GAPDH
CCL11
EEF2
SPP1
SOX9
PDL1
ACTB
GAPDH
CCL11
EEF2
SPP1
RUNX1
ACTB
PDL1
EEF2
GAPDH
RUNX1
SOX9
PDL1
ACTB
NANOG
COL1A1
GAPDH
EEF2
RUNX1
SPP1
NANOG
ACTB
GAPDH
CCL11
EEF2
COL1A1
SPP1
SOX9
RUNX1
ACTB
GAPDH
COL1A1
EEF2
SPP1
RUNX1
ACTB
GAPDH
COL1A1
EEF2
IL8
RUNX1
IL6
ACTB
GAPDH
CCL11
EEF2
COL1A1
SPP1
RUNX1
ACTB
PDL1
EEF2
GAPDH
RUNX1
ACTB
GAPDH
EEF2
SOX9
SPP1
RUNX1
ACTB
GAPDH
CCL11
EEF2
SPP1
RUNX1
ACTB
GAPDH
EEF2
RUNX1
PDL1
ACTB
NANOG
EEF2
GAPDH

**Figure S9. HUC showing heterogeneous spatial neighborhood networks**

Neighborhood network of a single HUC cell, revealing heterogeneity in different patches.

### HCH

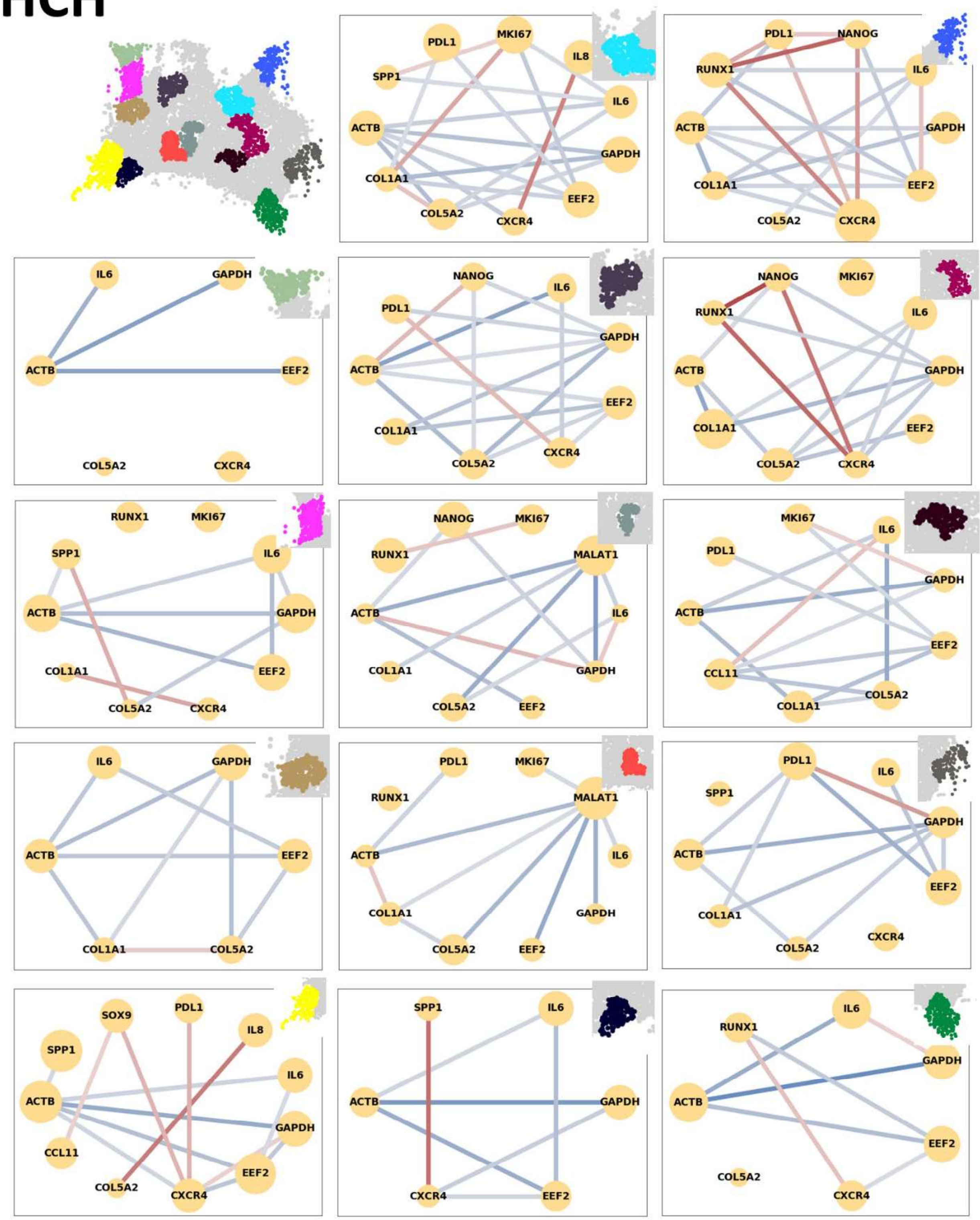
PDL1
MKI67
SPP1
IL8
IL6
ACTB
GAPDH
COL1A1
EEF2
COL5A2
CXCR4
PDL1
NANOG
RUNX1
IL6
ACTB
GAPDH
COL1A1
EEF2
COL5A2
CXCR4
IL6
GAPDH
ACTB
EEF2
COL5A2
CXCR4
NANOG
IL6
PDL1
GAPDH
ACTB
EEF2
COL1A1
CXCR4
COL5A2
NANOG
MKI67
RUNX1
IL6
ACTB
GAPDH
COL1A1
EEF2
COL5A2
CXCR4
RUNX1
MKI67
SPP1
IL6
ACTB
GAPDH
COL1A1
EEF2
COL5A2
CXCR4
NANOG
MKI67
RUNX1
MALAT1
ACTB
IL6
COL1A1
GAPDH
COL5A2
EEF2
MKI67
IL6
PDL1
GAPDH
ACTB
EEF2
CCL11
COL5A2
COL1A1
IL6
GAPDH
ACTB
EEF2
COL1A1
COL5A2
PDL1
MKI67
RUNX1
MALAT1
ACTB
IL6
COL1A1
GAPDH
COL5A2
EEF2
PDL1
IL6
SPP1
GAPDH
ACTB
EEF2
COL1A1
CXCR4
COL5A2
SOX9
PDL1
IL8
SPP1
IL6
ACTB
GAPDH
CCL11
EEF2
COL5A2
CXCR4
SPP1
IL6
ACTB
GAPDH
CXCR4
EEF2
IL6
RUNX1
GAPDH
ACTB
EEF2
COL5A2
CXCR4

**Figure S10. HCH showing heterogeneous spatial neighborhood networks**

Neighborhood network of a single HCH cell, revealing heterogeneity in different patches.

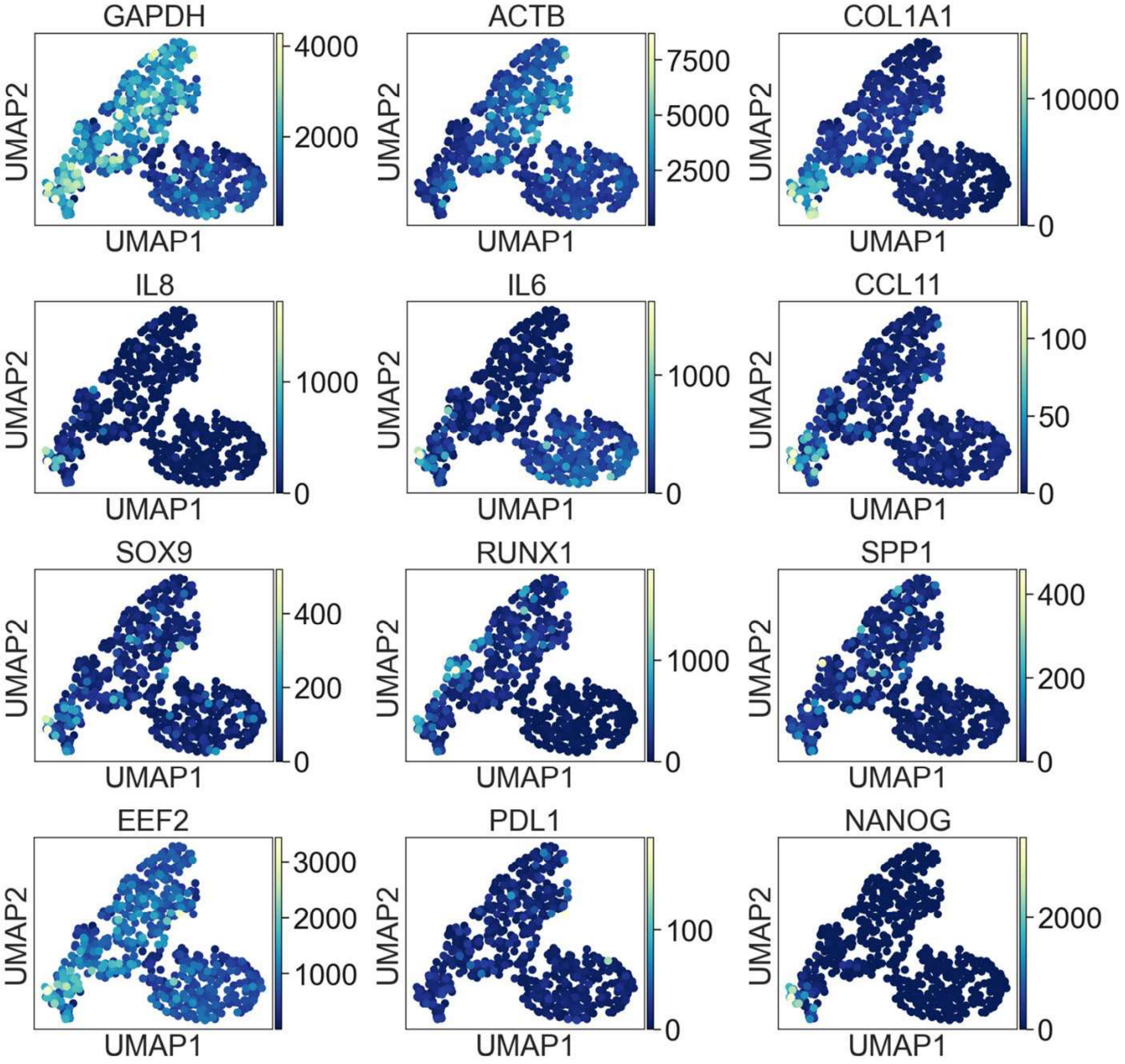
GAPDH
4000
2000
ACTB
7500
5000
2500
COL1A1
10000
0
IL8
1000
0
IL6
1000
0
CCL11
100
50
0
SOX9
400
200
0
RUNX1
1000
0
SPP1
400
200
0
EEF2
3000
2000
1000
PDL1
100
0
NANOG
2000
0
UMAP1
UMAP2

**Figure S11. Single-cell gene enrichment is associated with different regions on UMAP**

Single-cell RNA counts visualized in UMAP. Dots color indicates expression level of each gene of the cell.

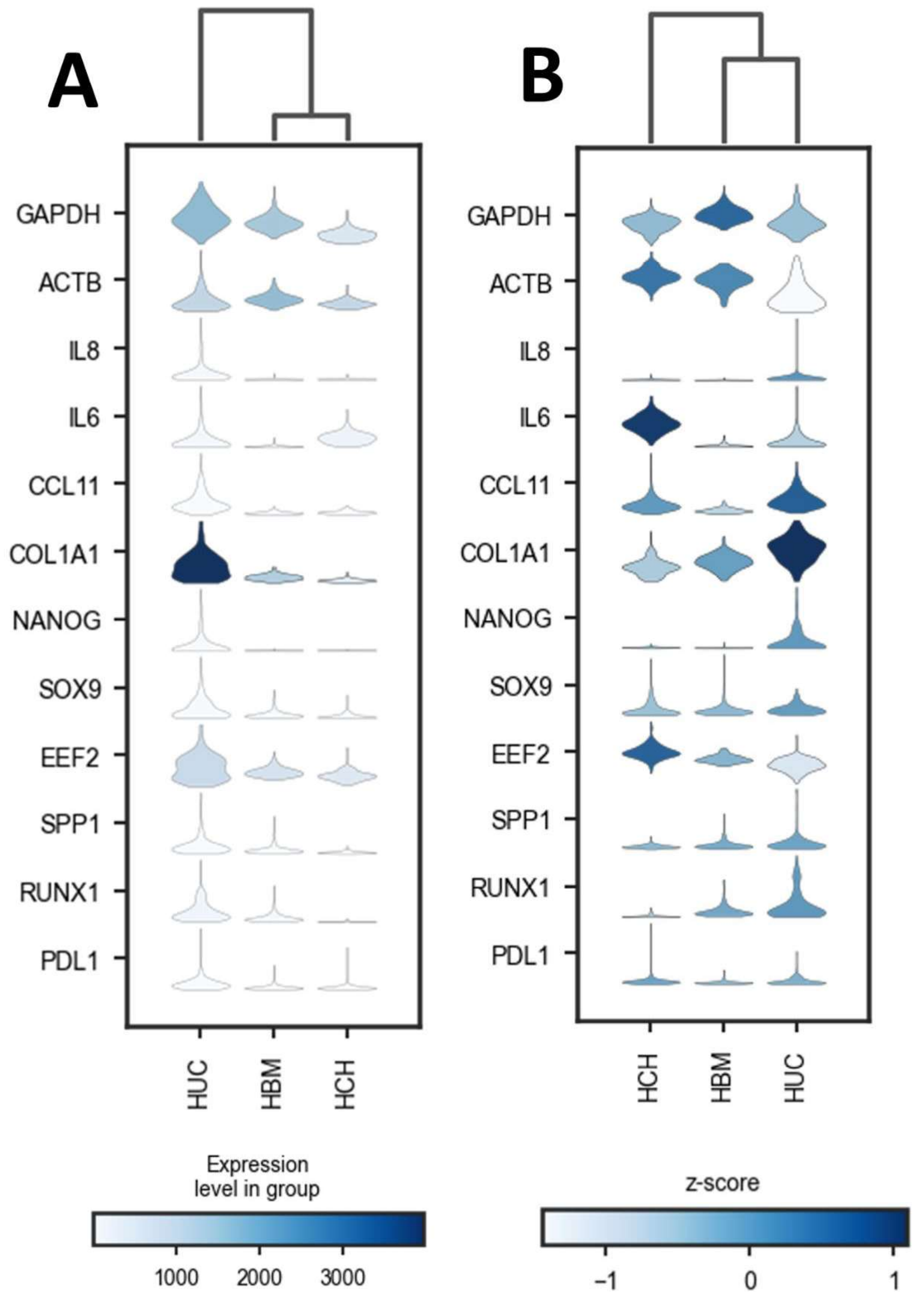
A
B
GAPDH
ACTB
IL8
IL6
CCL11
COL1A1
NANOG
SOX9
EEF2
SPP1
RUNX1
PDL1
HUC
HBM
HCH
HCH
HBM
HUC
Expression level in group
1000
2000
3000
z-score
−1
0
1

**Figure S12. Stacked violin plot of marker enrichment in each cell type**

(A) Stacked violin plot showing frequency of RNA-FISH gene counts of each cell type. The color of the violin plot indicates the average expression level of the marker in the cell type.

(B) Stacked violin plot shows the relative enrichment of markers in cell populations using the z-score of gene expression per cell. The color of the violin plot indicates the average z-score of the marker in each cell type.

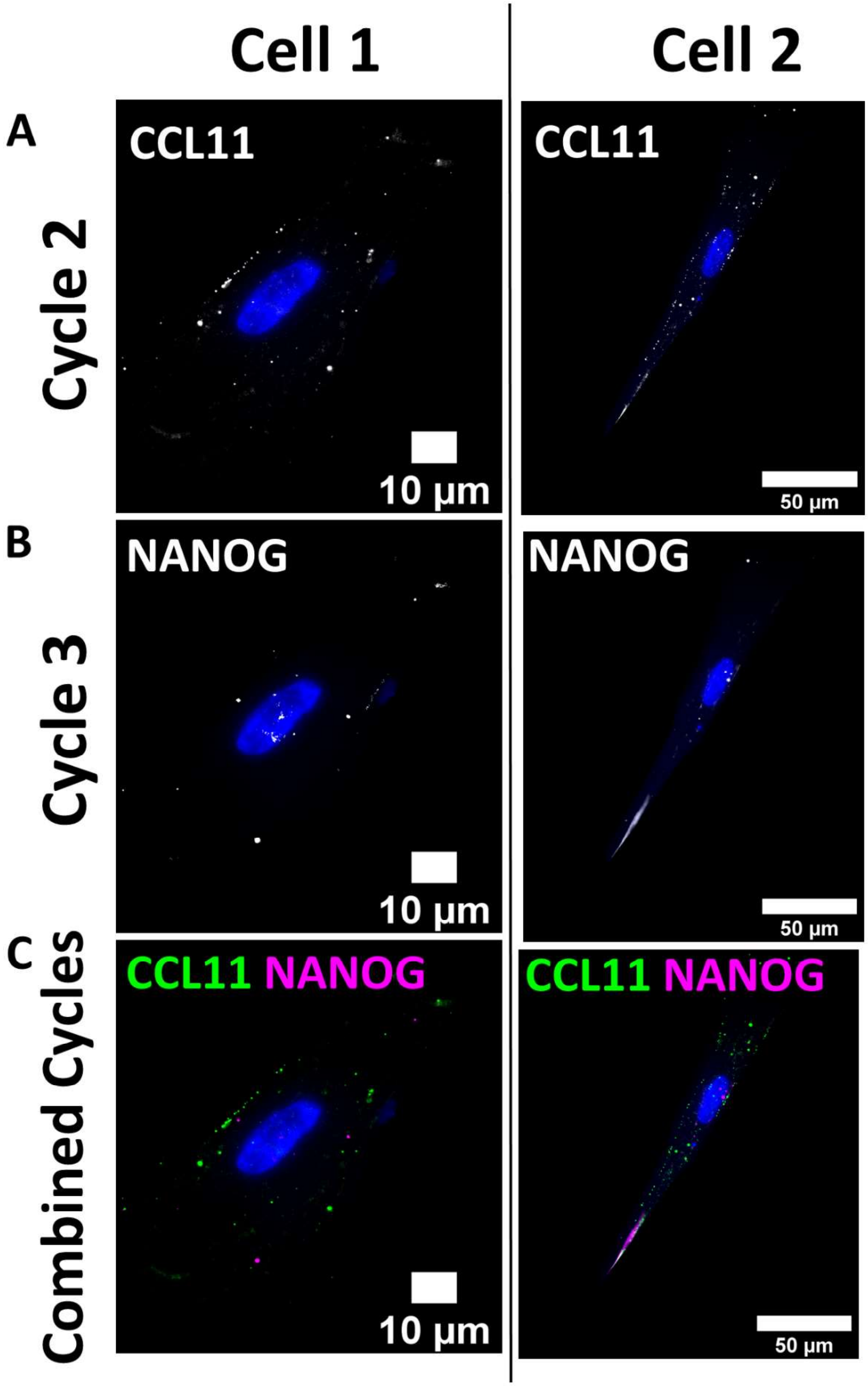

Cell 1
Cell 2
A
CCL11
CCL11
Cycle 2
10 µm
50 µm
B
NANOG
NANOG
Cycle 3
10 µm
50 µm
C
CCL11 NANOG
CCL11 NANOG
Combined Cycles
10 µm
50 µm

**Figure S13. Image of sequential cycles indicate removal of edge-localized cytokine RNA signal**

(A) Cy5 channel image of *CCL11*.

(B) Cy5 channel image of the following cycle. Gene labeled in Cy5 is *NANOG*.

(C) Comparison of Cy5 image of two sequential cycles. The change of fluorescent signal indicates the detected RNA signal is not residual staining.

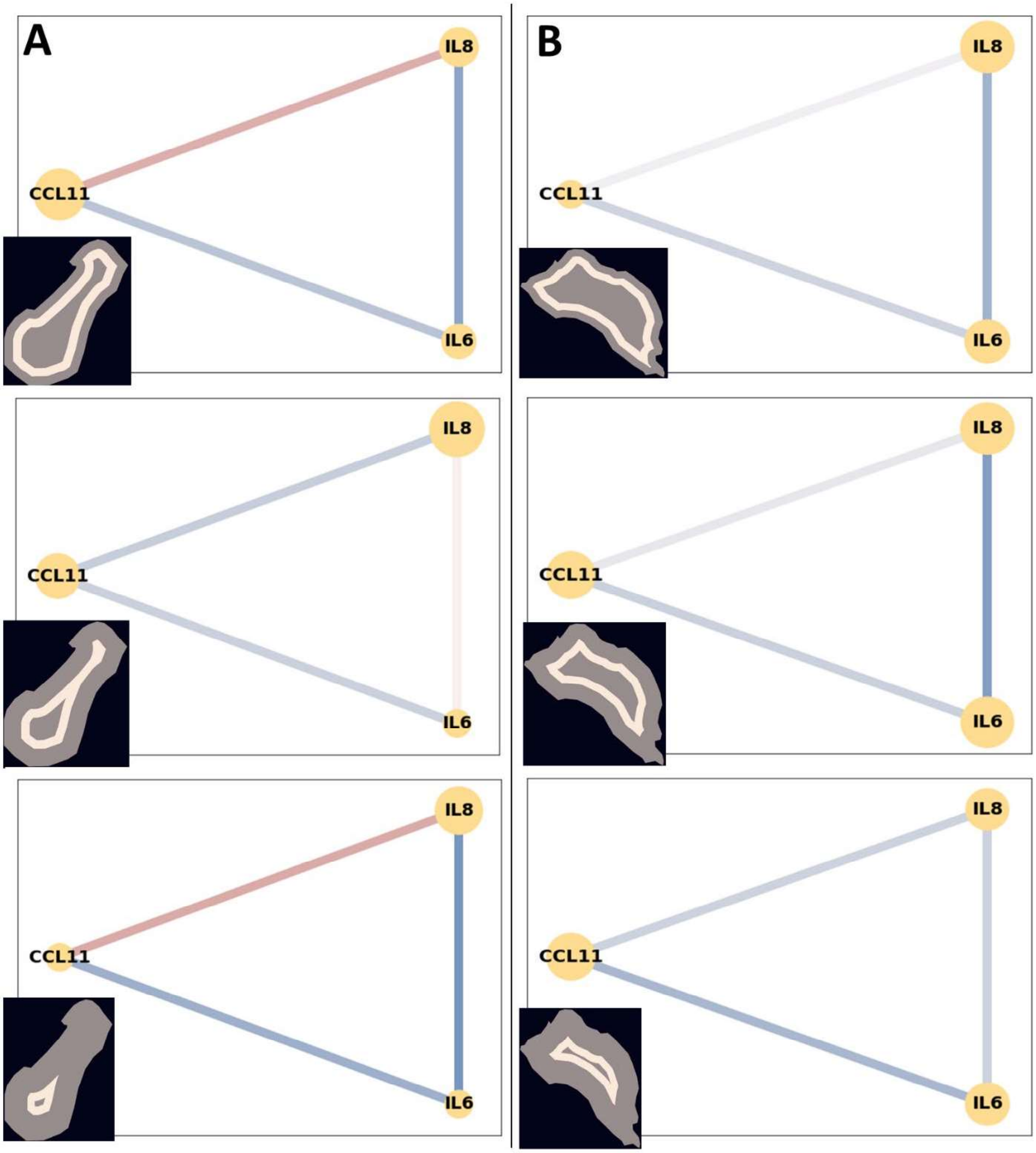
A
B
IL8
CCL11
IL6
IL8
CCL11
IL6
IL8
CCL11
IL6
IL8
CCL11
IL6
IL8
CCL11
IL6
IL8
CCL11
IL6

**Figure S14. Neighborhood networks of HUC show variability between different contour patches**

**(A)** Neighborhood networks of HUC with *CCL11* edge localization.

**(B)** Neighborhood networks of HUC without any edge localization.

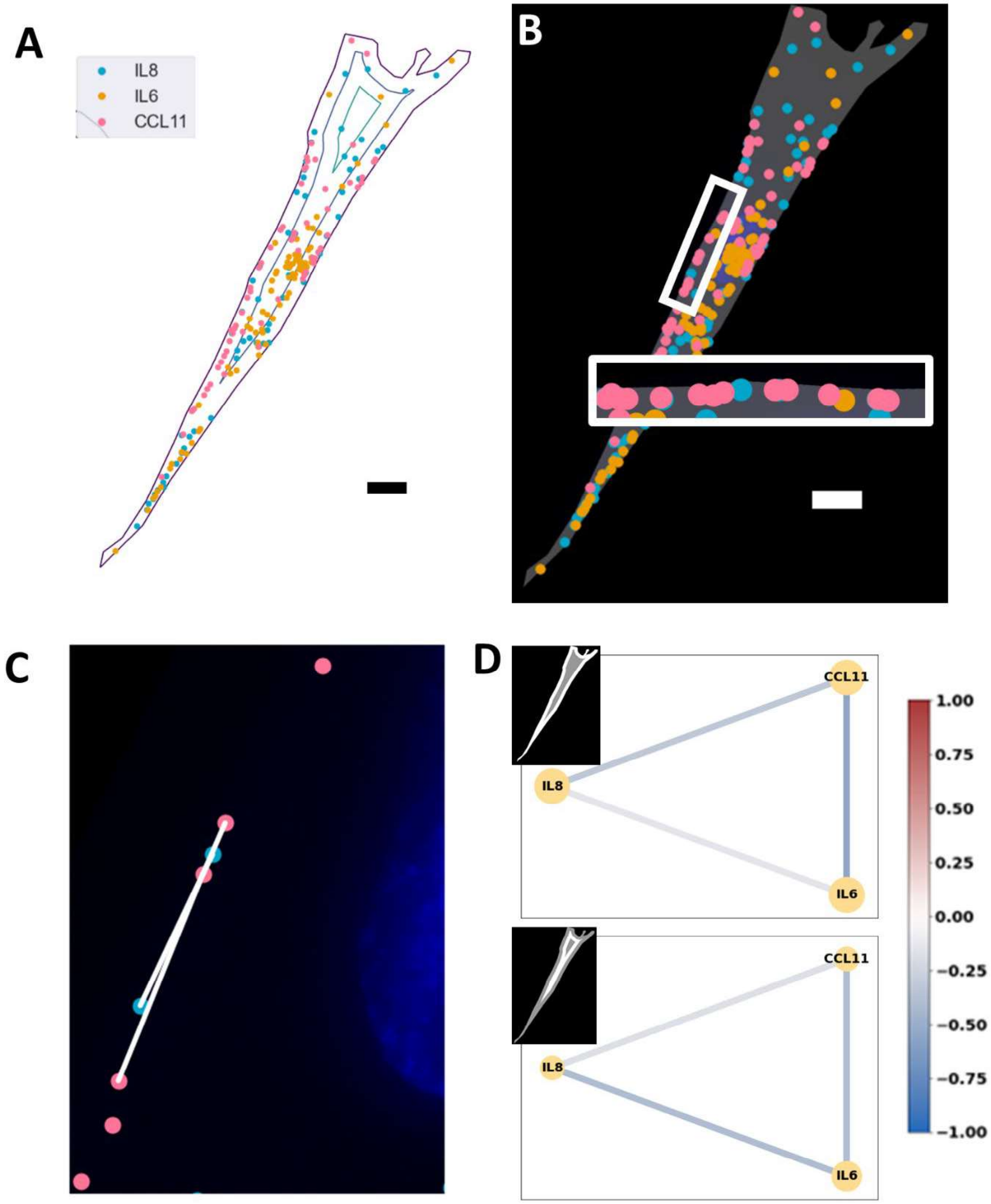
A
IL8
IL6
CCL11
B
C
D
CCL11
IL8
IL6
CCL11
IL8
IL6
1.00
0.75
0.50
0.25
0.00
−0.25
−0.50
−0.75
−1.00

**Figure S15. HUC with both *IL8* and *CCL11* edge-localization indicates different neighborhood networks**

**(A)** Contour patch map of HUC. Detected RNA are overlaid on contour plot. Gap between each equal distance line is a patch.

**(B)** Edge localization of *IL8* and *CCL11* RNA.

**(C)** A local neighborhood of the outermost patch. A center transcript and its 4 nearest neighbors form a local neighborhood. Connections follow the distribution of RNA along the edge of the cell.

**(D)** Neighborhood networks of contour patches. In this cell, *IL8* and *CCL11* show stronger negative correlation in the outermost patch. *IL6* and *IL8* show stronger negative correlation in the inner patch. Negative correlation between *IL6* and *CCL11* is preserved between the patches

Unlabeled scale bars have length of 10μm.

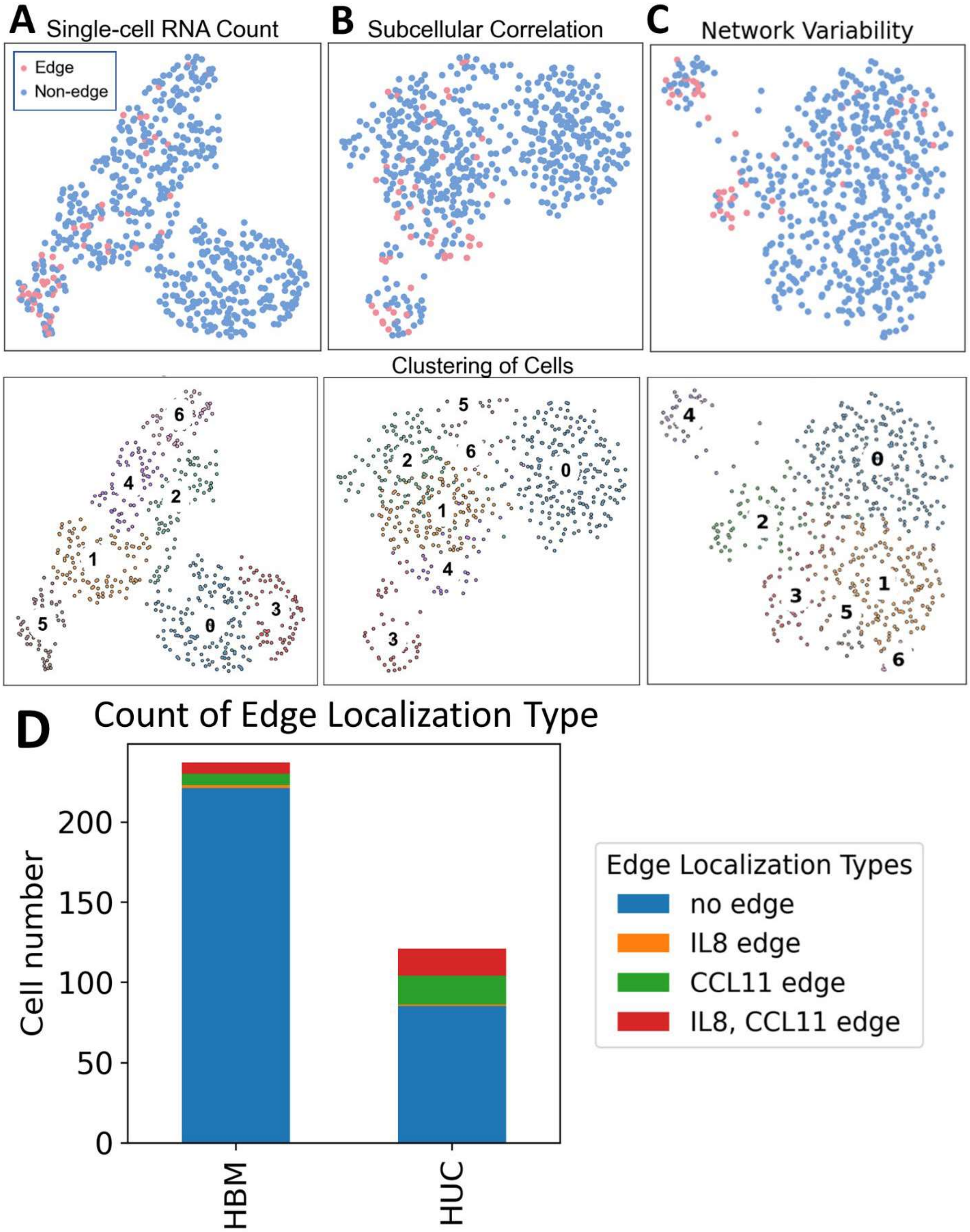

A
Single-cell RNA Count
Edge
Non-edge
B
Subcellular Correlation
C
Network Variability
Clustering of Cells
D
Count of Edge Localization Type
Cell number
0
50
100
150
200
HBM
HUC
Edge Localization Types
no edge
IL8 edge
CCL11 edge
IL8, CCL11 edge

**Figure S16. Visualization and statistics showing the proportion of MSCs with edge-localization of cytokine RNA.**

- **(A)** Cells with cytokine RNA edge-localization pattern highlighted in single-cell RNA count-based UMAP.
- **(B)** Cells with cytokine RNA edge-localization pattern highlighted in patch-level correlation-based UMAP.
- **(C)** Cells with cytokine RNA edge-localization pattern highlighted in neighborhood network variability based UMAP.
- **(D)** Proportion of HBMs and HUCs with edge-localized cytokine RNA.

| Cell | Source | Cat # | Age | Sex | Controls |
|---|---|---|---|---|---|
| MSC | Bone Marrow | #MSC-031<br>Lot#: 00238 | 22 | M | CD90+ CD166+<br>CD34- CD45-<br>C/A/O differentiation |
| MSC | Umbilical Cord | #C43001UC<br>Lot#: 210261 | N/A | F | CD90+ CD166+<br>CD34- CD45-<br>C/A/O differentiation |
| Chondrocyte | Hip | C-12710<br>Lot#: 445Z012.3 | 60 | M | Viability: 74%<br>PD Time: 66.3/PD |

**Table S1. Cells used for the study.**
